# Proximity interactome analysis of Lassa polymerase reveals eRF3a/GSPT1 as a druggable target for host directed antivirals

**DOI:** 10.1101/2021.07.16.452739

**Authors:** Jingru Fang, Colette Pietzsch, Haydar Witwit, George Tsaprailis, Gogce Crynen, Kelvin Frank Cho, Alice Y. Ting, Alexander Bukreyev, Erica Ollmann Saphire, Juan Carlos de la Torre

**Affiliations:** Department of Immunology and Microbiology, Scripps Research, La Jolla CA 92037; La Jolla Institute for Immunology, La Jolla CA 92037; Department of Pathology, University of Texas Medical Branch, Galveston, TX 77550; Galveston National Laboratory, University of Texas Medical Branch, Galveston, TX 77550; Proteomics Core, Scripps Research, Jupiter, FL 33458; Bioinformatics and Statistics core, Scripps Research, Jupiter, FL 33458; Cancer Biology Program, Stanford University, Stanford, CA 94305; Department of Genetics, Department of Biology and Department of Chemistry, Stanford University, Stanford, CA 94305; Chan Zuckerberg Biohub, San Francisco, CA 94158; Department of Microbiology and Immunology, University of Texas Medical Branch, Galveston, TX 77550

**Author notes:** Juan Carlos de la Torre. Erica Ollmann Saphire; **Email:**. **Author Contributions:** J.F., E.O.S., and J.C.T designed research; J.F. performed all non-BSL4 experiments and analyzed data; C.P. performed all BSL-4 experiments; A.B. provided supervision to BSL4 research; H.W. performed all LCMV experiments; G.T. performed proteomic experiments; G.C. performed the experimental design and data analysis for proteomics; K.F.C. and A.Y.T provided a critical resource; J.F., E.O.S., and J.C.T wrote the paper. All authors reviewed and edited the paper. **Competing Interest Statement:** The authors declare no competing interest.

**Keywords:** Viral replication, Proximity proteomics, Host-virus interactions

## Abstract

Completion of the Lassa virus (LASV) life cycle critically depends on the activities of the virally encoded RNA-dependent RNA polymerase in replication and transcription of the viral RNA genome in the cytoplasm of infected cells. The contribution of cellular proteins to these processes remains unclear. Here, we applied proximity proteomics to define the interactome of LASV polymerase in cells, under conditions that recreate LASV RNA synthesis. We engineered a LASV polymerase-biotin ligase (TurboID) fusion protein that retained polymerase activity and successfully biotinylated the proximal proteome, which allowed the identification of 42 high-confidence LASV polymerase interactors. We subsequently performed an siRNA screen to identify those interactors that have functional roles in authentic LASV infection. As proof-of-principle, we characterized eukaryotic peptide chain release factor subunit 3a (eRF3a/GSPT1), which we found to be a proviral factor that physically associates with LASV polymerase. Targeted degradation of GSPT1 by a small molecule drug candidate, CC-90009, resulted in strong inhibition of LASV infection in cultured cells. Our work demonstrates the feasibility of using proximity proteomics to illuminate and characterize yet to be defined, host-pathogen interactome, which can reveal new biology and uncover novel targets for the development of antivirals against highly pathogenic RNA viruses.

**Significance Statement:** Lassa virus (LASV), the causative agent of Lassa fever (LF), represents an important public health problem in Western Africa. There is no FDA-approved therapeutic intervention to treat LF. Due to its limited genome coding capacity, LASV proteins are often multifunctional and orchestrate complex interactions with cellular factors to execute steps required to complete the viral life cycle. LASV polymerase is essential for replication and expression of the viral genome, and thus is an attractive target for antiviral intervention. Here we present the first host interactome of the LASV polymerase, which can guide identification of novel druggable host cellular targets for the development of cost-effective antiviral therapies for LF.

## Introduction

Lassa virus (LASV) is a mammarenavirus that is highly prevalent in Western Africa, where it infects several hundred thousand individuals annually. A substantial number of LASV infections causes Lassa fever (LF), which has high morbidity and significant mortality among hospitalized LF patients (1). Although rodent-to-human is the main mode of transmission leading to LASV infections in human population, a significant number of LF cases can arise from human-to-human transmission (2). Moreover, increased travel has resulted in exported LF cases from endemic Western African countries to non-endemic countries (3). To date, no US Food and Drug Administration (FDA)-licensed countermeasures are available to prevent or treat LASV infections, and current anti-LASV therapy is limited to an off-label use of ribavirin that has limited efficacy and can cause significant side effects. Hence, there is an unmet need for cost-effective therapeutics to combat LASV infections. Development of effective therapeutics would be facilitated by a better understanding of virus-host cell interactions that modulate replication and gene expression of LASV in infected cells.

Like other mammarenaviruses (*Bunyavirales: Arenaviridae*), LASV is an enveloped virus with a bi-segmented, single-stranded RNA genome. Each viral genome segment uses an ambisense coding strategy to direct the synthesis of two viral proteins from open reading frames separated by non-coding intergenic region (IGR). The small (S) segment encodes the nucleoprotein (NP), which is responsible for genome encapsidation and immune evasion, and the glycoprotein precursor (GPC), which is co- and post-translationally processed to generate the mature GP that mediates virion cell entry via receptor-mediated endocytosis (4, 5). The large (L) segment encodes the large (L) protein that functions as a viral RNA-directed RNA polymerase (RdRp), and the matrix Z protein. NP encapsidates the viral genome and antigenome RNA species to form the viral nucleocapsid (NC) to which L associates to form the two viral ribonucleoprotein complexes (vRNPs) (of L and S segments). The resulting vRNPs are responsible for directing replication and transcription of the viral RNA genome (6, 7). Cellular polymerases and translation machinery can neither read nor translate the LASV RNA genome. Instead, the LASV polymerase initiates viral transcription from the genome promoter located at the 3’ end of the viral genome, which is primed by a small host-cell-derived, capped RNA fragment via a mechanism called cap-snatching. Primary transcription results in synthesis of NP and L mRNAs from the S and L segments, respectively. The virus polymerase can also adopt a replicase mode and moves across the intergenic region (IGR) to generate a copy of the full-length antigenome (cRNA) for each segment. These cRNAs serve as templates for synthesis of the GPC and Z mRNAs from the S and L segments, respectively, as well as templates for amplification of the corresponding viral genome (vRNA) (8–10).

Coordination of multiple domains with distinct enzymatic functions through conformational rearrangement and the engagement of vRNA promoter elements with LASV polymerase have been proposed to mediate the functional transitioning of LASV polymerase between a replicase and transcriptase (11). Here, we investigated whether host cell factors can critically contribute to the distinct steps of viral RNA synthesis that are driven by LASV polymerase. As a first and necessary step, we applied the recently developed proximity labeling technology to characterize the cellular interactome of LASV L polymerase in the context of viral RNA synthesis in living cells. We fused LASV polymerase to the engineered promiscuous biotin ligase TurboID to generate a fusion protein that retained polymerase activity and ability to biotinylate its cellular interactors *in situ*. This approach allowed direct affinity-capture and identification of biotinylated interactors, leading to the discovery of high-confidence, LASV polymerase interactors. To validate the functional role of the identified interactors, we implemented a high-content imaging-based, siRNA screen using a human hepatocyte derived cell line, Huh7 cells, infected with authentic LASV. Among the functional polymerase interactors, we identified GSPT1as a pro-viral factor of which pharmacological targeting with the pre-clinical drug CC-90009 had a strong inhibitory effect against LASV in cultured cells. Our results have improved our understanding of the cellular proteins and pathways involved in LASV RNA synthesis and uncover potential novel therapeutic strategies against a lethal human pathogen.

## Results

### Generating LASV L-HA-TurboID with polymerase and proximity labeling activities

For proximity labeling, we selected TurboID, an engineered biotin ligase that has promiscuous labeling activity and optimized labeling efficiency. TurboID labels exposed lysine residues on neighboring proteins within a ~10 nm radius and is highly active, requiring <10 minutes of labeling time for detectable activity (12). We incorporated TurboID with an HA tag into the LASV L protein at a site (after residue 407) previously shown to tolerate incorporation of epitope tags without severely affecting polymerase activity (13) (**Figure 1A**). The HA tag facilitated L-HA-TurboID detection and biochemical characterization as there is no commercially available antibody against LASV L.

**Figure 1.**
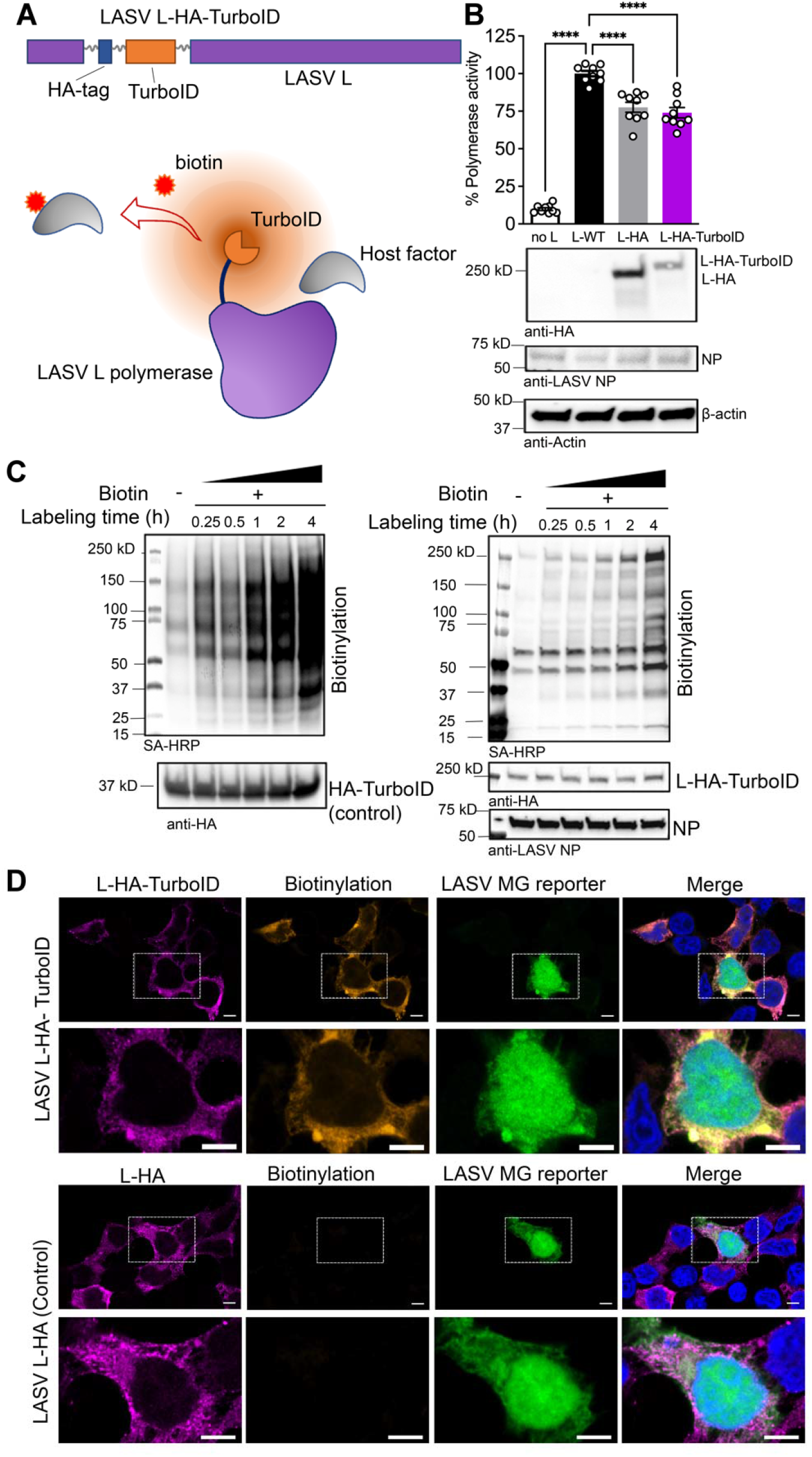
Generation of a functional LASV polymerase with proximity-labeling activity. **(A)** Schematic of LASV L-HA-TurboID construct and proximity biotinylation of L-interacting host factors. **(B)** Polymerase activity and expression of LASV L-TurboID compared to L-WT and L-HA in a cell-based LASV minigenome (MG) assay. MG activity (fluorescent reporter intensity) was quantified in each sample and normalized to the L-WT control. Representative western blots showing expression of LASV L-HA-TurboID, LASV L-HA, LASV NP and β-actin in HEK 293T whole cell lysates. MG activity was evaluated in three independent biological replicates with triplicated wells for each condition. Bar graphs containing nine individual data points (white circles) for each condition are displayed, with means plus standard error of the means/SEM (error bars). One-way ANOVA with Dunnett’s multiple comparisons test was performed to compare the activity of LASV L-HA-TurboID and L-HA to L-WT (****, *p* <□0.0001). **(C)** Streptavidin blot showing biotinylations mediated by a control HA-TurboID in the absence of LASV proteins (left panel) or by LASV L-HA-TurboID in cells co-expressing LASV MG components (right panel). **(D)** Confocal immunofluorescence images of LASV L-HA-TurboID mediated proximity biotinylation in cells co-expressing LASV MG components. LASV L-HA and L-TurboID were detected by using an anti-HA antibody. Biotinylated proteins were detected by using the streptavidin-AF594. Nuclei were stained with Hoechst and are shown in merged images. Scale bar: 5 μm. Representative images from two independent experiments acquired by Zeiss Laser-scanning microscopy (LSM)880 with Airyscan are shown. The lower row shows a zoomed-in view of the area outlined by a white dash line in the upper row. Each experiment includes at least three different fields of view.

We used a cell-based, LASV minigenome (MG) system to confirm the polymerase activity of the resulting L-HA-TurboID. This MG system recapitulates LASV RNA synthesis using an intracellular reconstituted LASV vRNP expressing a fluorescent reporter, ZsGreen. Reconstitution of LASV vRNP requires co-expression of the LASV L and NP proteins, as well as the LASV MG vRNA. Thus, we performed all experiments requiring reconstituted LASV MG in HEK 293T cells due to their high transfection efficiency. In these experiments, expression levels of the MG reporter served as a comprehensive measurement of LASV MG replication, transcription, and translation of the MG reporter transcript (14) (**Figure S1**).

We used a wild-type LASV L (L-WT) and an N-terminal HA tagged L (L-HA) as controls to compare the polymerase activity and protein expression levels with L-HA-TurboID. L-HA-TurboID retained 70% of WT activity, with slightly lower expression levels compared to that of the L-HA control (**Figure 1B**). We validated the proximity-labeling (PL) activity of L-HA-TurboID by detecting biotinylated proteins in cells reconstituted with LASV MG. In the presence of exogenous biotin, L-HA-TurboID produced a broad range of biotinylated proteins in a time-dependent manner. In comparison, a different group of cellular proteins was biotinylated by a N-terminal HA-tagged TurboID control (**Figure 1C**).

We next used confocal fluorescent microscopy to detect the subcellular distribution of L-HA-TurboID mediated biotinylation in cells. We transfected HEK 293T cells with a LASV MG system containing either L-HA-TurboID or L-HA control, and at 24 hours post-transfection conducted 1 hour biotin labeling. LASV L-HA-TurboID localized to discrete puncta connecting patches in the cytoplasm, similar to the L-HA control. Further, the biotinylation signal overlapped with the location of L-HA-TurboID, supporting efficient and localized biotinylation mediated by the L-HA-TurboID (**Figure 1D**).

### Defining LASV polymerase interactomes by proximity proteomics

We carried out a proximity proteomics experiment using similar conditions to those previously published (12). We transfected HEK 293T cells with the LASV MG system, including either L-WT or L-HA-TurboID followed by biotin labeling. Transfected cells were lysed, and biotinylated proteins were captured with streptavidin (SA) beads. Proteins enriched by SA beads were washed and subjected to on-bead trypsin digestion. The digested peptides in solution were then labeled with unique tandem-mass tags (TMTs) for quantitative proteomic analysis (**Figure 2A**).

**Figure 2.**
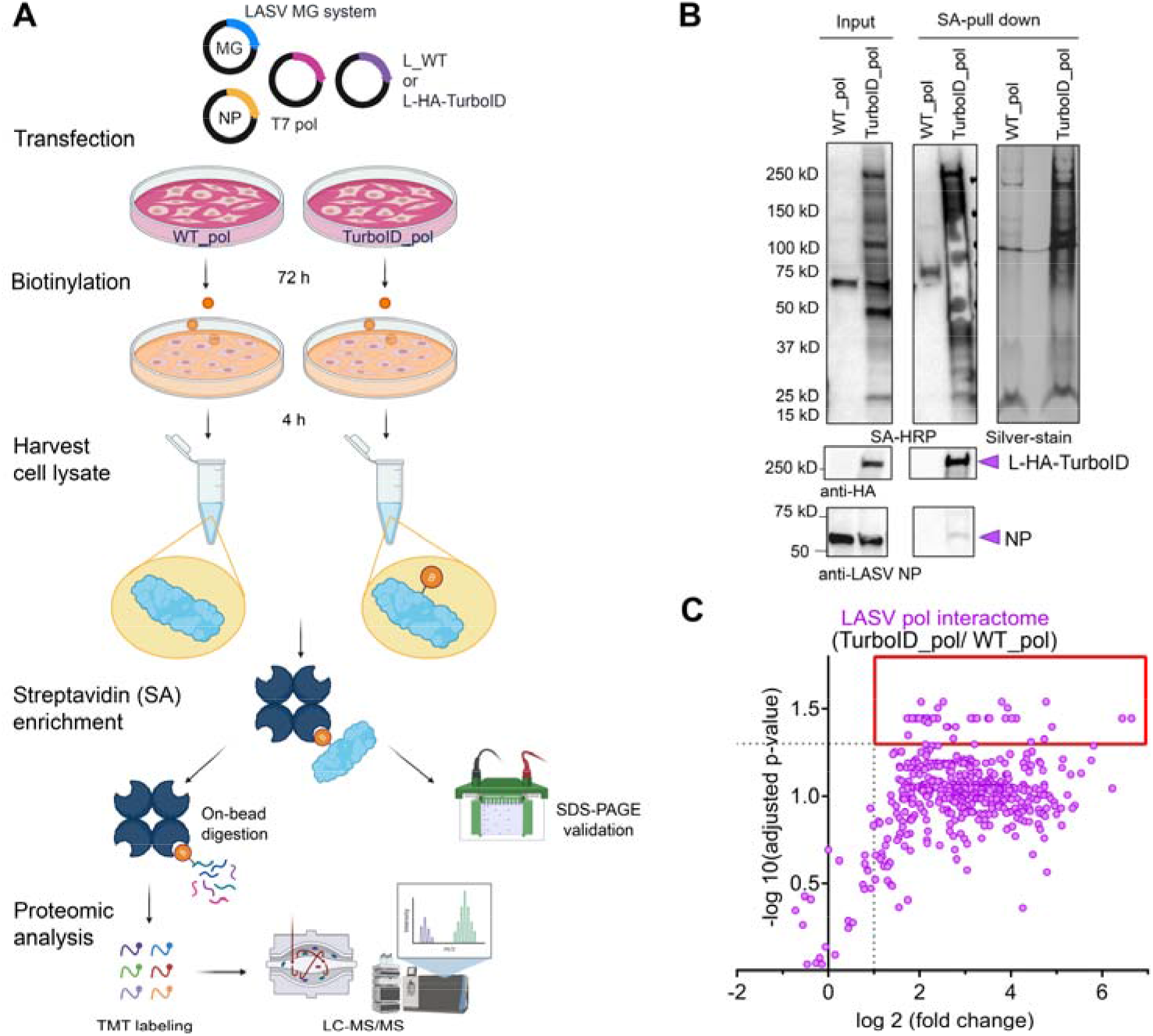
Proximity proteomics of LASV polymerase in living cells. **(A)** Sample preparation workflow for proximity proteomic assays. **(B)** Validation of streptavidin enrichment of LASV L-HA-TurboID mediated biotinylation. Harvested cell lysates are indicated as “Input” and biotinylated proteins enriched with streptavidin beads are indicated as “SA-pull down”. Representative blots and a silver-stained gel from three biological replicates are presented. **(C)** Scatter plot showing LASV polymerase interacting proteins in HEK 293T cells determined by quantitative proteomic analysis. The degree of enrichment for each protein is shown as a fold-change and the statistical confidence of enrichment is shown as adjusted p-values that are numerated and log-transformed. Data points within the red box corresponding to high-confidence hits.

To assess the efficiency of the enrichment, biotinylated material bound to SA beads (SA-pull down) from equal amounts of starting material (Input) was eluted using SDS loading buffer containing DTT and biotin and analyzed by western blot. We observed enrichment of biotinylated proteins specific to samples with L-HA-TurboID expression. Only a small fraction of LASV NP was biotinylated, which may reflect that only a low percentage of the total NP participates in the formation of a functional vRNP, or that in the vRNP the majority of NP was not accessible to biotinylation or remained insoluble under the lysis conditions we used to prepare the samples. We detected L-HA-TurboID in the SA-pull down fraction, suggesting self-biotinylation (**Figure 2B**).

To analyze the LASV polymerase interactome, host proteins identified as LASV polymerase interactors were scrutinized by two criteria: degree of enrichment compared to the control proteome, and the corresponding statistical confidence for each comparison. We used L-WT to generate the control proteome. For every identified protein, the abundance ratio in the L-HA-TurboID sample was normalized to that in the L-WT sample, to obtain a fold-change value. Multiple comparisons were performed across three biological replicates and adjusted P values (adjusted to False Discovery Rate of 0.05) were determined to identify proteins that were enriched in the L-HA-TurboID interactome. A threshold of adjusted *p* < 0.05 and log_2_(fold change) > 1 was used to identify 42 high-confidence hits corresponding to proteins that interact with LASV polymerase (**Figure 2C**).

### Effect of siRNA targeting high-confidence LASV polymerase interactors on viral infection

Next, we assessed the biological importance of the 42 high-confidence hits in the context of infection with live LASV. For this experiment, we used an siRNA-based functional screen based on well-established experimental procedures (15) and performed the screen using human hepatocyte-derived Huh7 cells, as hepatocytes are relevant to LASV induced pathology (16). Briefly, we transfected Huh7 cells with a library of individual siRNAs targeting each of the 42 high-confidence hits, and at 48 hours post-transfection infected these cells with the recombinant LASV-eGFP using a multiplicity of infection (MOI) of 0.5 (PFU/cell) (17). Using a high-content imaging system, we quantified the percentage of LASV-infected cells with each siRNA knockdown (KD) based on the number of GFP positive cells and the total cell count (with nuclear stain) (**Figure 3A**). The siRNA screen was repeated using a MOI of 1.

**Figure 3.**
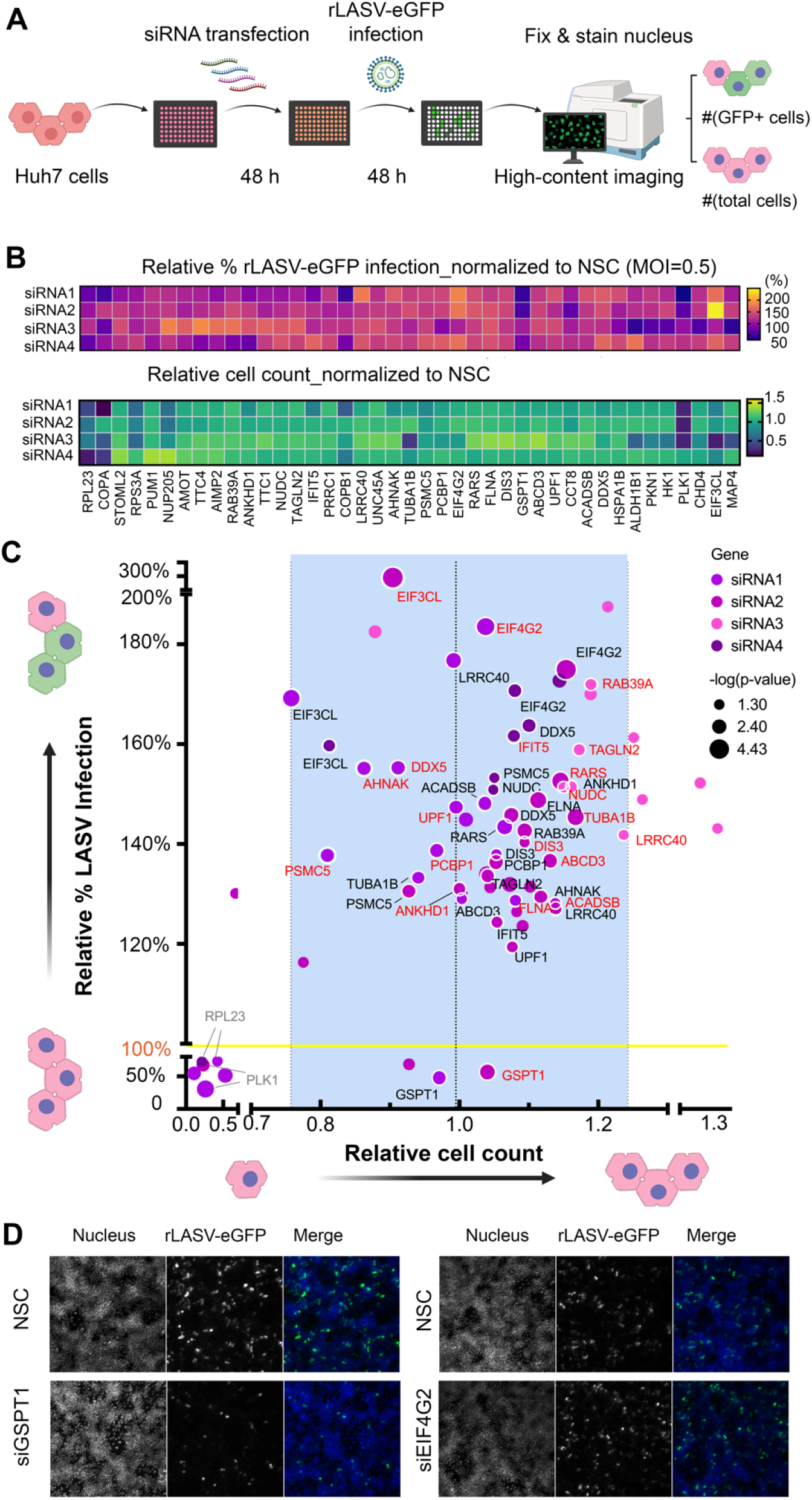
Identification of functional polymerase interactors by siRNA screen in LASV-infected cells. **(A)** Workflow of siRNA screen by high-content imaging of Huh7 cells infected with rLASV-eGFP. The raw percentage of infection and total cell count for each siRNA treatment were calculated and normalized to that of non-silencing controls (NSC). **(B)** Heat maps of relative percentage of infection and relative total cell count for LASV. Each value is the mean of technical triplicates. Multiple unpaired t-tests were performed to determine the statistical significance of siRNA-mediated changes in the percentage of infection compared to that for NSC. **(C)** The *p*-values were log-transformed and displayed as the size of each data point on a scatter plot. Data points in the heatmap with *p* > 0.05 were eliminated from the scatter plot. Names of genes for which multiple siRNAs significantly affected infection, but not cell counts, are highlighted on the corresponding data points. For each gene, only one label highlighted in red for display purposes. **(D)** Representative fluorescence images of Huh7 cell monolayers at 48 hours post infection are shown for NSC and two selected siRNAs. Merged images are composed of green (rLASV-eGFP) and blue (nuclei) channels. Results of the siRNA screen with MOI=0.5 is shown.

To control potential non-specific effects of siRNA transfection on LASV infection and total cell count, we normalized these two parameters against values for non-silencing control (NSC) siRNA. We excluded any hits for which siRNA-KD altered the total cell count by one standard-deviation from the average cell count or for which siRNA-KD failed to significantly change the percentage of LASV-infected cells compared to NSC. From the remainder, we considered as true hits those with at least two independent siRNA-KDs resulted in the same infection phenotype.

19 high-confidence hits exhibited an antiviral role, as depletion of the hit by more than two independent siRNAs increased LASV infection (MOI=0.5) (**Figure 3B**). Six of these antiviral hits were validated in the repeated siRNA screen (MOI=1) (**Figure S2**). Limited (10 - 20% of cells) infection of NSC siRNA transfected cells (**Figure S3**) may have favored the detection of enhancement over reduction in infection. Top hits in the siRNA screen were selected by stringent statistical metrics (two biological replicates, four independent siRNAs per target, technical triplicate per siRNA) and were guided by quantitative parameters of the resulting phenotype (i.e., relative % infection and cell count). These top hits, included six antiviral factors (EIF3CL, EIF4G2, TAGLN2, TUBA1B, PSMC5 and UPF1) and one pro-viral factor (GSPT1).

### The proximity interactome of LASV L polymerase

We summarized the LASV L polymerase interactome in a network representation, in which we highlighted hits that were functionally validated in our siRNA screen. We clustered all interactors based on their STRING-functional classification (18) to identify distinct cellular pathways relevant to LASV infection (**Figure 4A**).

**Figure 4.**
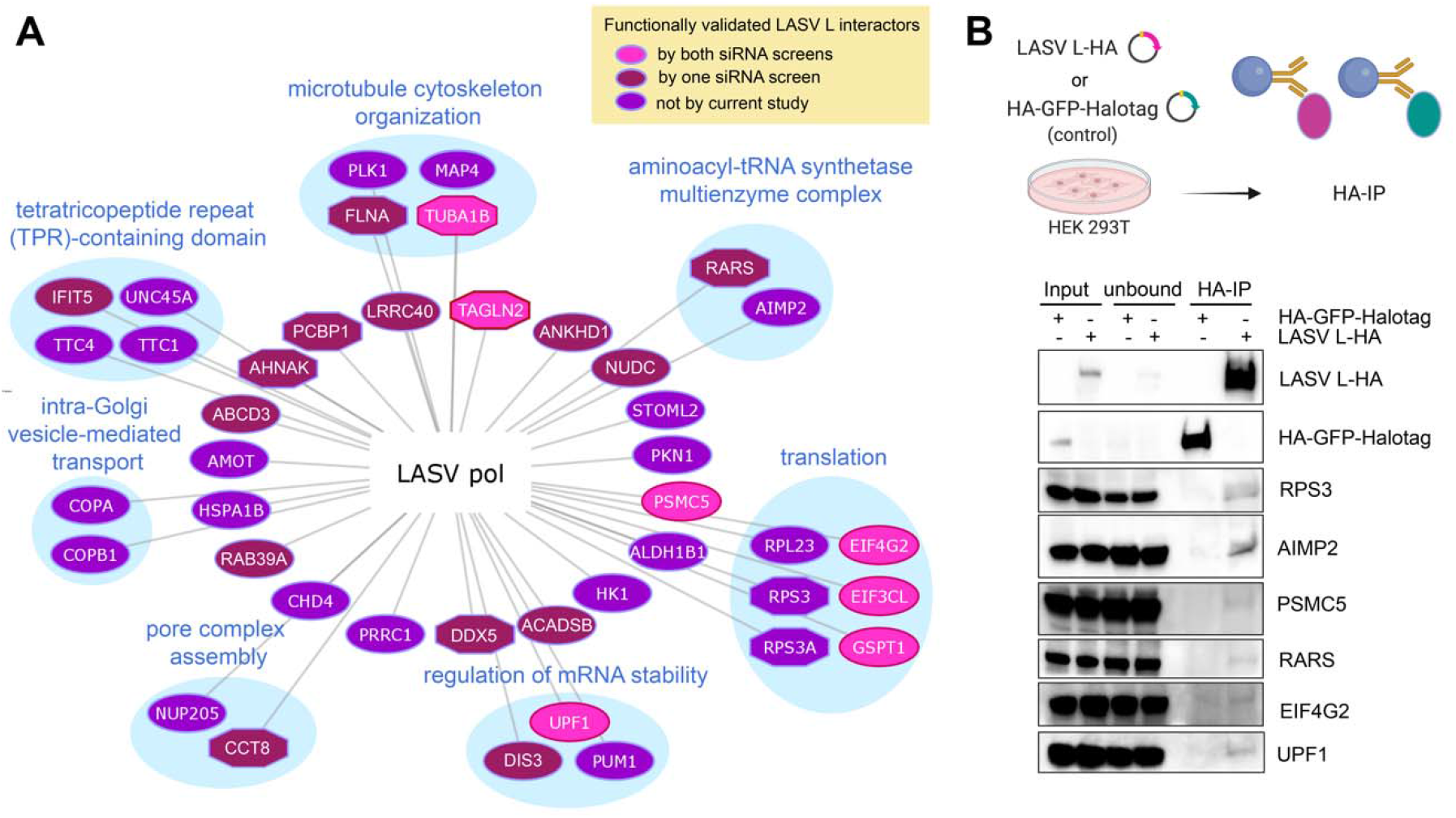
The proximity interactome of LASV L polymerase. **(A)** Network of the LASV polymerase interactome identified by proximity proteomics. Each node represents one high-confidence proteomic hit. Each edge represents an identified protein-protein interaction. Nodes that were functionally validated by siRNA screen are highlighted in pink (by two siRNA screens), or in maroon (by one siRNA screen). Nodes shown in heptagons have been previously reported for other virus-host interactomes. Selected nodes are clustered based on enriched biological processes or protein domains by functional enrichment analyses via STRING. **(B)** Validation of selected proteomic hits by co-immunoprecipitation of HA-tagged LASV L transiently expressed in HEK 293T cells. HA-tagged GFP-Halotag fusion protein was used as a control. Representative blots of two experiments are shown.

To confirm the LASV L interactome, we selected eight interactors for biochemical validation. We coimmunoprecipitated (co-IP) endogenous LASV L interacting proteins in HEK 293T cells using an N-terminal HA-tagged L as the bait (**Figure 4B**). Among the 8 selected interactors, we validated RARS, AIMP2, RPS3, PSMC5, EIF4G2 and UPF1. GSPT1 and AMOT were the two interactors that we could not successfully co-IP, likely due to the low expression levels of the corresponding endogenous proteins. To control non-specific bindings, we performed a parallel co-IP using an HA-tagged GFP-Halotag fusion protein as the bait. These results support the validity of proximity labeling-based proteomics to identify *bona fide* LASV L-interactors.

We also compared the LASV L interactome described here with previously reported host interactomes of viral proteins derived from the prototypic Old World (OW) arenavirus, lymphocytic choriomeningitis virus (LCMV) (**Figure S4**). We found 11 common host factors shared by multiple interactomes, which likely reflects conserved molecular recognition interfaces between host cells and OW mammarenaviruses.

### Role of GSPT1 in LASV gene expression

As proof-of-principle, we focused on the only proviral hit, eukaryotic peptide chain release factor subunit 3a (eRF3a/GSPT1), which has not been reported to play any specific role in viral infection.

First, we used co-IP to verify that GSPT1 physically interacts with LASV L protein. To ensure the level of GSPT1 protein was sufficient for co-IP, we co-transfected HEK 293T cells with plasmids expressing an N-terminal Flag-tagged GSPT1 (long isoform, 68.7 kDa) and LASV L-HA, in the presence or absence of other LASV MG components. We found that LASV L-HA consistently immunoprecipitated Flag-GSPT1 in the presence or absence of LASV NP, or LASV NP and the LASV minigenome (**Figure 5A**). We next examined the functional consequence of GSPT1 association with LASV L by the LASV MG assay. We found that depletion of GSPT1 protein by one experimentally validated siRNA, GSPT1si8, (**Figure S5**) significantly suppressed LASV MG activity, but did not affect expression of a control eGFP reporter (**Figure 5B**).

**Figure 5.**
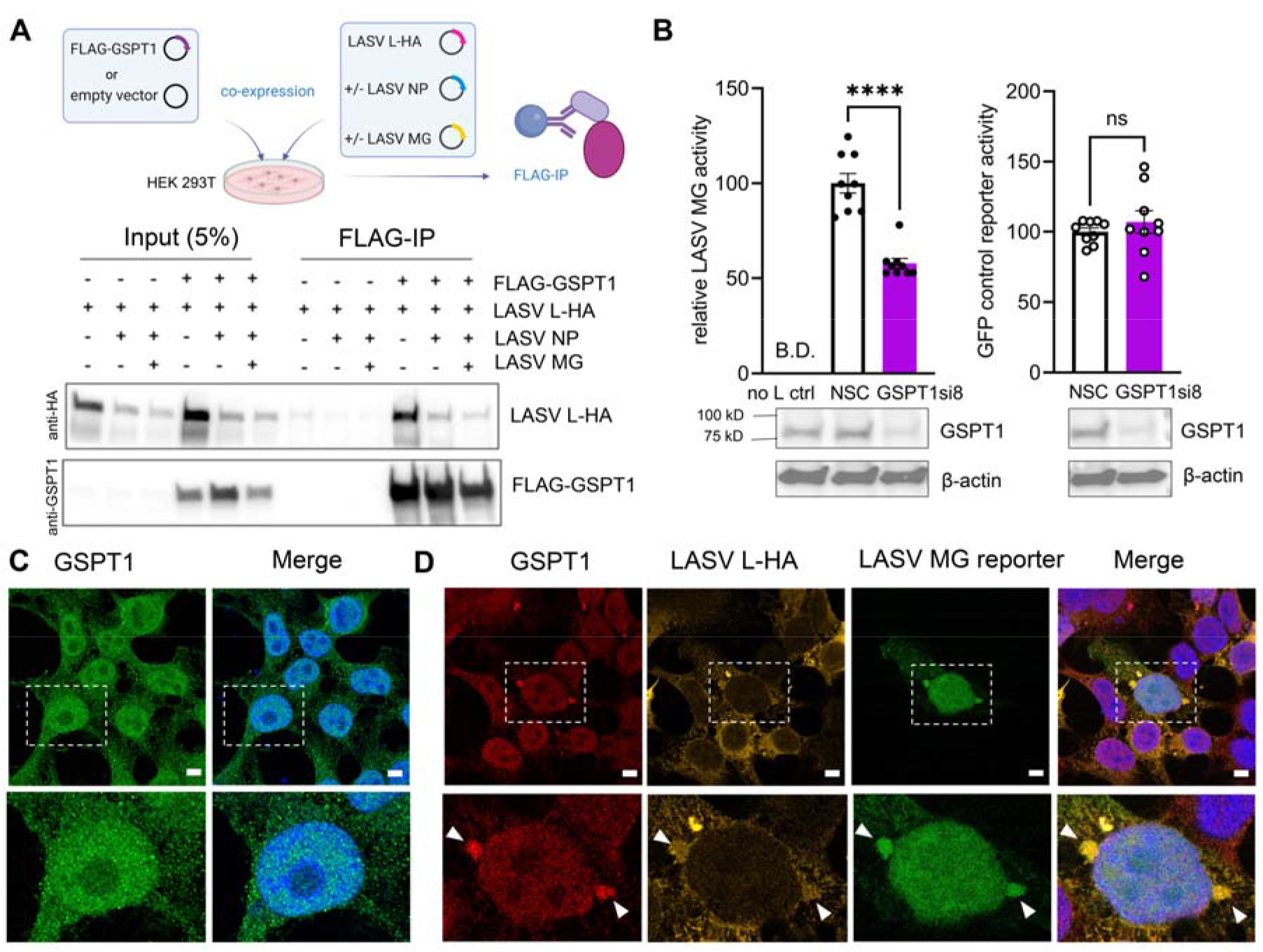
GSPT1 physically and functionally associates with LASV polymerase. **(A)** Western blot analyses of FLAG-IP from cells co-expressing FLAG-GSPT1 and LASV L-HA with or without LASV minigenome (MG) components. Representative results from three biological replicates are shown. **(B)** LASV MG activity and expression level of a control GFP reporter in HEK 293T cells upon GSPT1-knockdown (KD). GSPT1-KD by siRNA was validated by western blot analysis. β-actin served as the loading control. MG activity and expression level of the control GFP reporter in GSPT1si8-transfected cells were normalized to that of NSC transfected cells. The LASV MG experiment was repeated in three independent replicates with triplicate wells for each condition. Nine individual data points for each condition are displayed in the bar graphs, with means ± SEM (error bars) shown. Two-tailed, unpaired t tests were performed to determine whether depleting GSPT1 significantly changed activities of LASV MG and the control reporter (*ns: not significant;* ****, *p* <□0.0001). Confocal immuno-fluorescent analysis of the subcellular localization of endogenous GSPT1 alone **(C)** or with transiently expressed LASV L-HA in LASV MG reconstituted HEK 293T cells **(D)**. Nuclei were stained with Hoechst (blue) and are shown in merged images. Arrowheads: sites where endogenous GSPT1 localize proximal to LASV proteins. Representative images from two independent experiments that were acquired with Zeiss-LSM880 Airyscan are shown, each experiment includes at least three different fields of view. The lower row shows a zoomed-in view of the area outlined by a white dash line in the upper row. Scale bar: 5 μm.

We also examined whether endogenous GSPT1 interacts with LASV polymerase in a cellular context that recapitulates viral RNA synthesis. In HEK 293T control cells, endogenous GSPT1 exhibited a diffuse nucleocytoplasmic distribution (**Figure 5C**). However, in HEK 293T cells that successfully reconstituted a functional LASV vRNP, as determined by the ZsGreen reporter signal, we detected a fraction of cytoplasmic GSPT1 that co-clustered with the signal for LASV L-HA (**Figure 5D**), suggesting that GSPT1 interacts with LASV L in cells expressing a functional LASV vRNP.

### Role of GSPT1 in LASV multiplication

To validate the role of GSPT1 on LASV multiplication, we examined the phenotype of siRNA-mediated GSPT1-KD in multi-step LASV growth kinetics in Huh7 cells. Consistent with the siRNA screen results, GSPT1 depletion significantly impaired LASV growth in Huh7 cells starting at 24 hours post-infection (h.p.i.) with at least one log reduction in viral titer at 48 h.p.i. (**Figure 6A**). The phenotype of impaired LASV growth in GSPT1-KD cells was observed in three biological replicates (**Figure 6A** and **Figure S6**). Upon GSPT1-KD, none of the LASV RNA species were altered at 6 h.p.i. (**Figure 6B**), whereas at 72 h.p.i. the level of LASV c/mRNA levels were reduced (**Figure S6B**). Notably, in GPST1-depleted cells at 72 h.p.i. LASV GP2 protein levels were decreased, which might impede production of infectious virions (**Figure 6C** and **Figure SC**). This finding can be related to GSPT1 being a critical component of the cellular translation termination complex (eRF1/eRF3) (19–21). Given the subtle impact on LASV RNA accumulation but decrease in LASV protein levels, we reason that could GSPT1 supports LASV infection by assisting viral protein translation.

**Figure 6.**
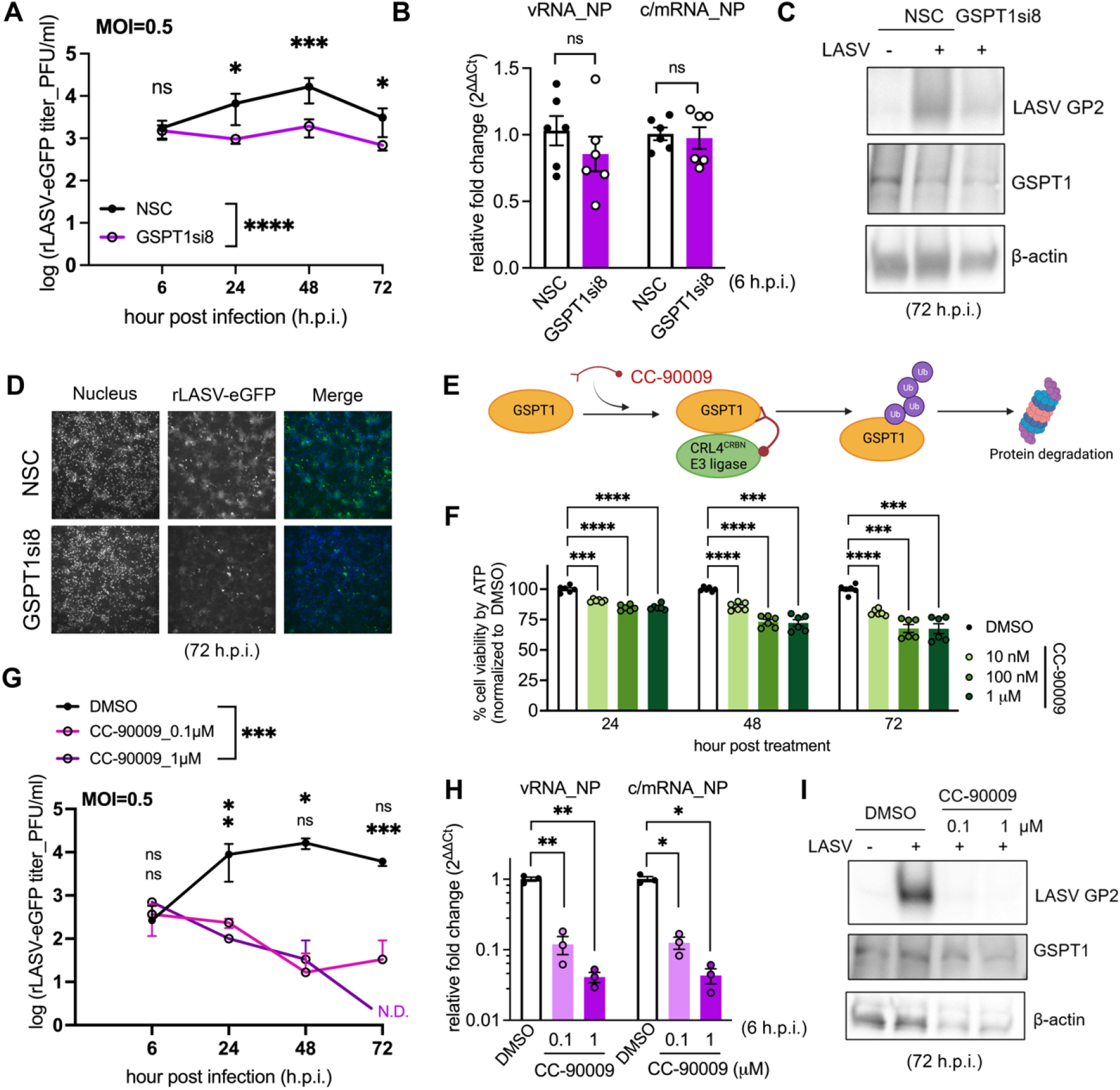
GSPT1 is a druggable host factor required for LASV multiplication. **(A)** Effect of GSPT1-knockdown (KD) on LASV viral growth kinetics in Huh7 cells. Viral titers from two independent experiments with technical triplicates were log-transformed and plotted as mean ± SD (error bars). Two-way ANOVA analysis with Šídák’s multiple comparisons test on log-transformed titers was performed to determine the statistical significance of the effect of GSPT1-KD on viral growth kinetics and on viral titers at each time point, respectively (*ns, not significant*; *, *p* <□0.05; **, *p* <□0.01; ***, *p* <□0.001; ****, *p*<0.0001). **(B)** Effect of GSPT1-KD on LASV vRNA and c/mRNA accumulation. Six individual data points from two independent experiments with technical triplicates are displayed and the mean ± SEM (error bars) is shown. Welch’s t test was performed to determine the statistical significance of the effect of GSPT1-KD on LASV RNA accumulation. **(C)** Western blot analysis of endogenous GSPT1 protein and LASV GP2 levels in lysates from LASV-infected Huh7 cells. β-actin served as the loading control. Lysates from triplicate wells in the growth kinetic experiments were pooled and analyzed. **(D)** Representative fluorescence images of Huh7 monolayers at 72 hours post-infection (h.p.i.) are shown with nuclei (blue) and virus infected cells (green). Representative blots and images from one experiment are shown. **(E)** Schematic diagram CC-90009-mediated GSPT1 degradation. **(F)** Cell viability of Huh7 cells treated with CC-90009 in the absence of viral infection was determined by Cell titer Glo 2.0. Raw values for cell viability in drug-treated samples are normalized to a DMSO control. Six individual data points from two independent experiments with technical triplicates are combined and indicated as mean ± SEM (error bars). Values that differed significantly from the controls were determined by two-way ANOVA with Dunnett’s multiple comparisons test. **(G)** Effect of CC-90009 on LASV growth kinetics in Huh7 cells. Results of one experiment with technical triplicates were plotted as mean ± SD (error bars). N.D: not detectable. Two-way ANOVA analyses on log-transformed viral titers with Dunnett’s multiple comparisons test were performed to determine the statistical significance of the effect of CC-90009 treatments compared to DMSO on LASV growth kinetics and viral titers at each time point, respectively. **(H)** Effect of CC-90009 treatment on LASV vRNA and c/mRNA accumulation in Huh7 cells. Three individual data points from one experiment with technical triplicates are displayed as mean ± SEM (error bars). Brown-Forsythe and Welch ANOVA with multiple comparisons test were used to determine values that differed significantly from the controls. **(I)** Western blot analysis of endogenous GSPT1 protein and LASV GP2 levels in LASV-infected Huh7 lysates. Lysates from triplicate wells in the growth kinetic experiments were pooled and analyzed.

Based on the effect of GSPT1 knockdown on LASV growth kinetics, we predicted that pharmacological inhibition of GSPT1 would suppress LASV multiplication. We therefore tested the antiviral activity of CC-90009, an E3 ubiquitin ligase modulator, which selectively tethers GSPT1 to the CRL4^CRBN^ E3 ubiquitin ligase to induce targeted ubiquitination and degradation of GSPT1 (22, 23) (**Figure 6D**). We treated Huh7 cells with CC-90009 immediately following LASV infection and measured the effect of CC-90009 treatment on LASV growth kinetics, viral RNAs, and viral protein accumulation. At 0.1 and 1 μM, CC-90009 effectively inhibited LASV growth (**Figure 6E**), which correlated with the dose-dependent reduction in accumulation of both vRNA and c/mRNAs accumulation within 6 hours of viral infection (**Figure 6F**) and diminished viral protein accumulation compared to the DMSO-treated control (**Figure 6G**). Importantly, in the absence of viral infection, we did not observe severe cytotoxicity of CC-90009 (**Figure 6H**), or reduction in cellular β-actin levels in the treated Huh7 cells (**Figure S5B**), indicating that the growth defect of LASV was highly unlikely caused by cytotoxicity of CC-90009 treatment.

We noticed that, unlike GSPT1-depletion mediated by siRNA, CC-90009 treatment equivalently inhibited accumulation of both LASV vRNA and c/mRNA at very early after infection (compare **Figure 6H** to **6B**), suggesting a fundamental defect in LASV RNA synthesis. This finding, together with the physical association between GSPT1 and LASV L polymerase, infer that CC-90009tethering of the E3 ligase to GSPT1 might facilitate ubiquitination and degradation of LASV L. On the other hand, as CC-90009 being able to modulate the spectrum of substrates targeted by the CRL4^CRBN^ E3 ligase (24, 25), LASV L itself could be turned into a neosubstrate and targeted for degradation upon CC-90009 treatment. Thus, reduced level of LASV L could entail a global decrease in LASV RNA synthesis, although this hypothesis remains to be examined. Together, our results demonstrate that CC-90009 has potent antiviral activity against LASV in human hepatocytes by targeting GSPT1.

### Effect of GSPT1 depletion on other mammarenaviruses

To determine whether the antiviral activity of CC-90009 could be expanded to other Old World (OW) mammarenaviruses, we examined the effect of CC-90009 on the growth kinetics of lymphocytic choriomeningitis virus (LCMV), an OW mammarenavirus related to LASV, in Huh7 cells. Delayed LCMV growth was seen in Huh7 cells infected with LCMV and then treated with CC-90009 (0.1μM) following infection, and with 1 μM CC-90009 LCMV titers were sustainably reduced by more than one log (**Figure 7A**). Similar to the phenotype observed in LASV-infected cells, accumulation of LCMV RNAs (**Figure 7B**) and proteins (**Figure 7C**) was reduced in CC-90009 treated cells, and fewer cells were infected with LCMV compared to vehicle treated cells (**Figure 7D**).

**Figure 7.**
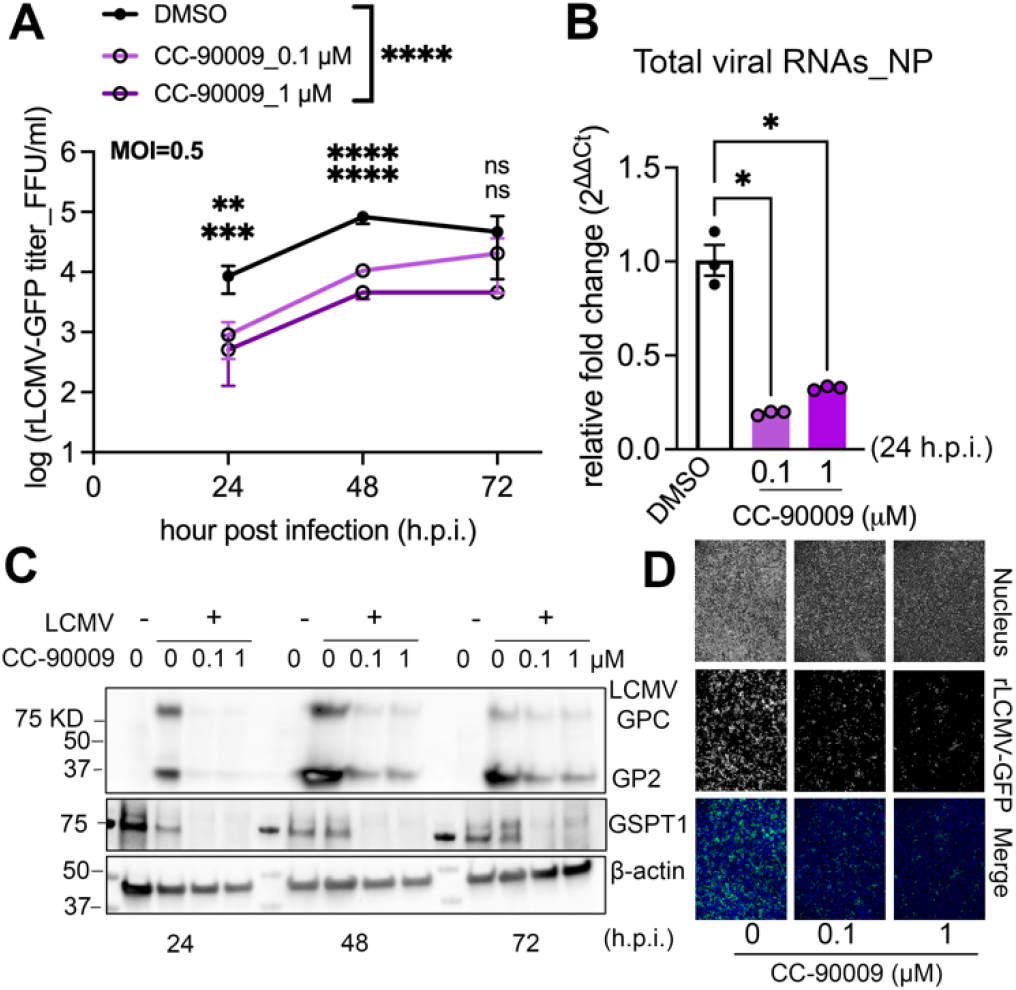
Inhibition of LCMV growth by CC-90009 mediated degradation of GSPT1. **(A)** Effect of CC-90009 on LCMV growth kinetics in Huh7 cells. Results of two independent experiments with technical duplicates are plotted as mean ± SD (error bars). Two-way ANOVA analyses of log-transformed viral titers with Dunnett’s multiple comparisons test were performed to determine the statistical significance of the effect of CC-90009 treatments compared to DMSO on LCMV growth kinetics and viral titers at each time point, respectively. **(B)** Effect of CC-90009 treatment on LCMV RNA synthesis in Huh7 cells. Three individual data points from one experiment with technical triplicates are displayed and indicated as mean ± SEM (error bars). Values significantly different from the controls were determined by Brown-Forsythe and Welch ANOVA with multiple comparisons test. **(C)** Western blot analysis of endogenous GSPT1 protein, LCMV GPC and GP2 levels in LCMV-infected Huh7 lysates. β-actin was used as the loading control. Lysates from duplicated wells in the growth kinetic experiments were pooled and analyzed. (**D)** Representative fluorescence images of Huh7 monolayers infected with rLCMV-GFP at 72 hours post-infection (h.p.i.) are shown with nuclei (blue) and virus-infected cells (green). Representative blots and images from one experiment are shown.

## Discussion

Here we have presented the first host interactome of LASV polymerase in the context of viral RNA synthesis directed by an intracellularly reconstituted functional LASV vRNP. We applied proximity proteomics to identify the LASV polymerase interactors *in situ*. We then performed an siRNA screen in human hepatocytes infected with live LASV to identify polymerase interactors functionally important during LASV infection. Our finding that most identified polymerase-interactors exhibited an antiviral rather than proviral phenotype could reflect that the viral polymerase or viral RNA products, or both, are targets of host innate defense mechanisms. Nevertheless, we validated one proviral factor, GSPT1, and demonstrated that it is a potentially druggable target for the development of therapeutics against LASV.

Previously reported host interactomes on LCMV proteins were based on affinity purification-mass spectrometry (AP-MS) approach in the context of viral infection (26, 27), which may fail to capture non-stable transient protein-protein interactions. Moreover, the use of live LASV infection requires BSL4 containment, which complicates proteomic experiments using LASV-infected cells. To overcome these obstacles, we used the TurboID-based, proximity-labeling approach to capture both stable and dynamic protein-protein interactions in the context of a cell-based MG system that recapitulates the biological activities of the LASV vRNP. Detection of endogenous host proteins expressed at low levels can be difficult to capture in AP-MS based interactomes. In proximity proteomics, nevertheless, direct enrichment of biotinylated interactors to the bait protein offers a higher likelihood of detecting host factors that have low abundance yet important roles such as GSPT1. This technological advantage may have contributed to the identification of 31 novel cellular proteins not identified by the LCMV L-interactome.

Many RNA viruses have evolved strategies to manipulate the host translational machinery to favor translation initiation of viral mRNAs (28–30). We identified two translation initiation factors in our LASV polymerase interactome, EIF4G2 and EIF3CL, as repressors of LASV infection. EIF4G2 is a functional homolog of EIF4G1 (31), which is known to bridge the high-affinity cytoplasmic cap-binding protein EIF4E and the scaffolding protein EIF3, which binds to the small ribosomal subunit. EIF4G2 can also enhance the interaction between EIF4E and cap-containing mRNAs (32). As with other mammarenaviruses, transcription of LASV mRNAs uses a ‘cap-snatching’ mechanism by which viral mRNAs hijack 5’ cap structures from host mRNA species (33), which primes viral transcription and enables cap-dependent translation (34). Although still controversial, one mechanism by which mammarenaviruses could accomplish “cap-snatching” involves binding of virus NP to m7GpppN cap structures (35, 36), together with the activity of an endonuclease motif (EndoN) present within the N-terminal region of the LASV polymerase (37) that cleaves cellular mRNAs to liberate the 5’ cap structure. Importantly, our LASV L interactome included EIF4G2 and EIF3CL, but not EIF4E, suggesting that LASV polymerase may not hijack the canonical host cap-binding complex. Our results could indicate that L polymerase instead competes for 5’ capped cellular mRNAs with the canonical host cap-binding complex, which contains EIF4E, and L may bind to EIF4G2 as a decoy that interferes with stable complex formation between eIF4E and capped mRNAs. This binding, in turn, would release capped mRNAs that can then serve as substrates for the L EndoN to hijack the 5’ cap structure for LASV transcription initiation. Additional support for this model stems from the observation that cap-binding affinity of EIF4E was reduced upon binding of LASV Z protein to EIF4E (38), which further lowers the affinity barrier for a LASV encoded cap-binding protein, or another pro-viral factor, to capture fragments of capped cellular mRNAs for priming LASV transcription (**Figure S7**).

Our siRNA-based functional screen also revealed LASV polymerase interactor, ATP-dependent RNA helicase upstream frameshift 1 (UPF1) as an antiviral factor. UPF1 is a key player in multiple cellular RNA decay pathways, typically through binding to the 3’ UTR of a target transcript (39), suggesting that UPF1 might play an antiviral role in LASV infection by mediating LASV mRNA decay. LASV mRNAs could trigger UPF1-dependent mRNA decay through either the nonsense-mediated mRNA decay (NMD) pathway or another mechanism resembling the replication-dependent histone mRNA decay pathway (HMD) (**Figure S8**). In the NMD pathway, UPF1 degrades aberrant RNA transcripts by joining the translation-termination complex (GSPT1/eRF3-eRF1) upon ribosomal recognition of a premature termination codon (PTC) in the mRNA (40). After recruitment to the termination complex, UPF1 is phosphorylated, thereby triggering a downstream cascade of nucleases and de-capping enzymes to dismantle the target transcript (41). LASV mRNAs are likely to incorporate premature termination codons owing to the presence of an upstream open reading frame (uORF), which can be acquired by viral transcripts as byproducts of the cap-snatching mechanism (42). In the HMD pathway, UPF1 phosphorylation is associated with recognition of the 3’ UTR stem-loop structure on histone mRNA by the step-loop-binding protein (SLBP). Similar to histone mRNAs, which lack 3’ poly(A) tail, LASV mRNAs also are not poly-adenylated and harbor a stem-loop structure at the 3’ end (43, 44) due to structure-dependent transcription termination at the LASV intergentic region (IGR) (45, 46). Future experiments are needed to determine whether the UPF1-LASV L interaction is RNA-dependent, and whether UPF1 directly targets LASV mRNAs for degradation. SLBP was not captured in the LASV polymerase interactome, but we cannot rule out the possibility that SLBP recognizes the 3’ stem-loop on LASV mRNAs. These findings provide the first evidence of a possible participation of the host RNA quality control mechanisms on modulating mammarenavirus infection.

Intriguingly, we identified another host NMD factor, G1-to-S-phase transition 1(GSPT1)/eukaryotic peptide chain release factor GTP-binding subunit A(eRF3a) as a LASV L interactor with pro-viral activity. GSPT1 is a core component of the cellular translation termination machinery (20, 47), has been shown to regulate cell cycle progression (48) and promote apoptosis (49). As GSPT1 participates in multiple cellular pathways, it may regulate viral infection via different mechanisms in a context-dependent manner. We confirmed a physical association between LASV L polymerase and GSPT1. Further, we showed that GSPT1 positively regulates LASV MG activity, arguing against the role of GSPT1 in the NMD being linked to its pro-viral activity. In the context of a multi-step growth kinetics experiment, GSPT1 knock-down resulted in one-log reduction of the number of infectious LASV progeny as well as reduced levels of both viral RNA and protein. These results point to a supporting role of GSPT1 in LASV gene expression, which could reflect a critical dependency of LASV on cellular translation termination machinery for its propagation in host cells. Whether the pro-viral phenotype we observed for GSPT1 is indirectly mediated by its involvement in regulating the cell cycle or apoptosis remains to be determined. Future studies are needed to resolve the functional consequence of the physical association between GSPT1 and LASV polymerase in LASV gene expression and to verify whether GSPT1 contributes to any step of LASV RNA synthesis independent from its potential impact on LASV protein translation.

CC-90009, a CRL4^CRBN^ E3 ligase modulating drug that specifically targets GSPT1 for proteasomal degradation (22), exhibited antiviral activity against both LASV and LCMV without appreciable cytotoxicity. For both viruses, CC-90009 treatment reduced accumulation of viral RNA and protein in infected cells, suggesting that similar mechanisms drive CC-90009 antiviral activity against LASV and LCMV. Based on the physical interaction between LASV L and GSPT1, it is plausible that LASV L protein might have a higher turnover rate upon CC-90009 treatment. However, due to the lack of access to an antibody to LASV L, we were not able to directly monitor L protein in our viral growth curve experiment. Further validation of this hypothesis can be enabled by monitoring the turnover rate of an epitope or reporter tagged LASV L in CC-90009 treated cells in the presence or absence of GSPT1. CC-90009 seemed to have a greater inhibitory effect on LASV than LCMV in Huh7 cells. However, future experiments should determine if GSPT1 also interacts with LCMV L and investigate the different sensitivity of LCMV infection to CC-90009-mediated inhibition. Lastly, whether CC-90009 also inhibit any of the human pathogenic New World mammarenaviruses remains to be examined. The fact that CC-90009 is currently in phase 1 clinical development for the treatment of acute myeloid leukemia raises the possibility of its repurposing as an antiviral drug against LASV and LCMV.

There are several limitations in the current work to be discussed. First, the use of wild-type LASV L as the control in our proximity proteomic experiment does not control for non-specific interactions associated with the TurboID tag in the LASV L-HA-TurboID fusion protein. This might have resulted in some false positives in our list of proteomic hits. However, a control proximity proteome generated by TurboID alone could have resulted in the elimination of cellular proteins that interact with LASV L, because both LASV L and TurboID are cytoplasmic proteins that have no clear physical boundary that would constrain their localizations. To support the specificity of our interactome, we biochemically validated a subset of the identified interactors via co-IP using an HA-tagged L that lacks TurboID. We confirmed that seven out eight (87%) tested hits interacted with LASV L. Moreover, we utilized the siRNA-based functional screen to support the relevance of LASV L interactome. We confirmed 21 functional hits among 42 interactors (50%) in at least one siRNA screen, including seven top hits in two independent siRNA screens using different infection conditions. It is possible that some of LASV L interactors we identified in HEK 293T cells are cell-type specific, whereas we performed our siRNA screen in Huh7 cells. We used HEK 293T cells in several experiments because their high transfection efficiency, and Huh7 cells in all viral infection experiments, as liver is a major target organ during LASV infection. Future works should confirm the functional role of LASV L interactors in other primary target cells including macrophage and dendritic cells, as well as in suitable animal models. Taken together, we presented here the first cellular interactome of LASV L polymerase, which illuminated new mechanisms of host regulation of LASV replication and revealed a landscape of novel targets for host-directed antiviral drug development.

## Materials and Methods

### Plasmids, siRNAs, antibodies, primers, and compound

see SI Appendix, Table S1, Dataset S2, Table S2, and Table S3.

### Cell cultures and viruses

see SI Appendix.

### LASV minigenome assay

see SI Appendix.

### Biotinylation with TurboID fusion proteins

See *SI Appendix.*

### Immunofluorescent analysis (IFA) using confocal microscopy

see *SI Appendix*.

### Proximity labeling-based proteomics (sample preparation, quality control, proteomic experiment, and data analysis)

See *SI Appendix.*

### siRNA functional screening and data analysis

Huh7 cells were transfected with individual gene-targeting siRNA and infected with rLASV-eGFP (MOI=0.5 or 1). Percentage of LASV infected cells and total cell counts were quantified at 48 hours post infection. See *SI Appendix* for details.

### LASV growth kinetics experiment

Huh7 cells were transfected with siRNA targeting GSPT1 and infected with rLASV-eGFP (MOI=0.5) 48 hours post-transfection; or were infected with rLASV-eGFP (MOI=0.5) and treated with CC-90009 at the specified concentration one hour post infection. At indicated time points, LASV vRNAs or c/mRNAs, proteins, and titers were quantified. See *SI Appendix* for details.

### LCMV growth kinetics experiment

rLCMV/GFP infected Huh7 cells were treated with CC-90009 at the specified concentration one hour post infection. At indicated time points, LCMV RNAs, proteins, and titers were quantified. See *SI Appendix* for details.

### Co-immunoprecipitation

See *SI Appendix*.

### RT-qPCR

See *SI Appendix.*

### Cell viability quantification by Cell-titer Glo assay

See *SI Appendix.*

## Acknowledgments

We thank Beatrice Cubitt from the de la Torre Lab (Scripps Research, CA) for helping with the cloning of pCI-FLAG-GSPT1 plasmid and preparation of LASV minigenome plasmid stocks. We thank Sharon Schendel at La Jolla Institute for Immunology (LJI) for manuscript editing, Zbigniew Mikulski of the Microscopy Core Facility (LJI) for microscopy training, and NIH S10OD021831 for sponsoring the Zeiss LSM 880 microscope. We thank Paul Schimmel (Scripps Research, CA) for providing us with anti-RARS and anti-AIMP2 antibodies, and Tianying Zhang (Scripps Research, CA) for the mouse monoclonal antibody to LCMV GP2. This research was supported by institutional funds of LJI (E.O.S.) and NIH/NIAID grants AI125626 and AI128556 (J.C.T.). J.F. was supported by the Donald E. and Delia B. Baxter Foundation Fellowship. This is publication # 30107 from Scripps Research.

