## Supplementary material for "Proximity interactome analysis of Lassa polymerase reveals eRF3a/GSPT1 as a druggable target for host directed antivirals": SI Appendix

**This PDF file includes:**

Figures S1 to S7
Legends for Datasets S1 to S2
Supplementary Methods
Tables S1 to S3
SI References

**Other supplementary materials for this manuscript include the following:**

Datasets S1
Datasets S2

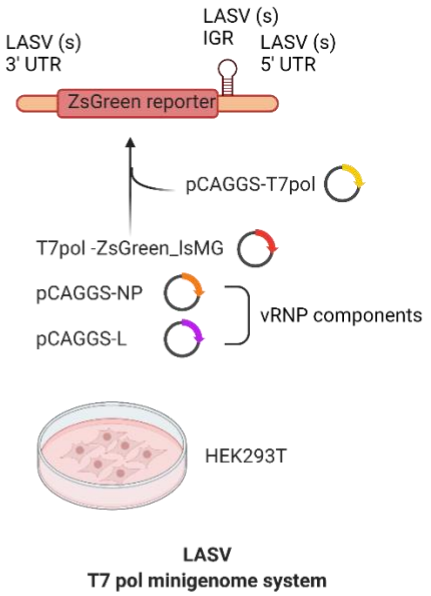


**Fig. S1. Schematic diagram of the LASV minigenome system.** Transfection of T7 RNA polymerase expression plasmid together with a plasmid encoding the ZsGreen reporter flanked by the 3′-and 5'-termini of the authentic LASV S genome segment and containing the corresponding intergenic region (IGR). T7 polymerase-dependent intracellular production of a reporter LASV minigenome (MG) can be subsequently replicated and transcribed by co-expressed LASV L polymerase and NP.

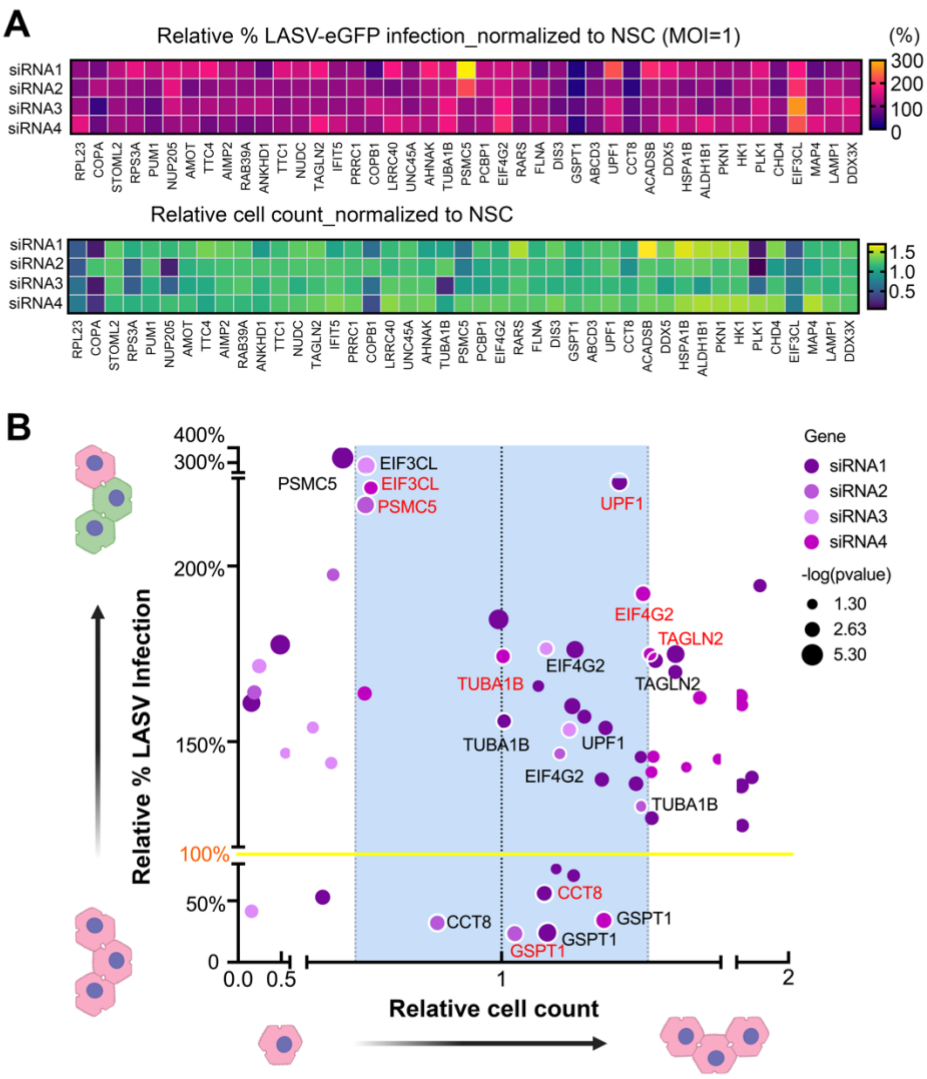


**Fig. S2. Relative percentages of viral infection and cell counts determined in siRNA-transfected cells. (A)** Raw percentage of infection and cell count for each siRNA treatment were normalized to that of non-silencing controls (NSCs). Normalized values are displayed in heat maps displaying relative percentage of infection and relative cell count for LASV. Each value is the mean of technical triplicates. Multiple unpaired t-tests were performed to determine the statistical significance of siRNA-mediated changes on the percentage of infection compared to that of NSC. **(B)** Calculated *p-values* were log-transformed and displayed as the size of each data point on a bubble plot. Data points in the %infection heat map with values exceeding the plotted range are marked bright yellow. **(C)** Data points in the heatmap with a *p-value* >0.05 are not shown in the bubble plot. Names of genes for which multiple siRNAs significantly affected infection are labeled on the corresponding data points; For each gene, only one label is marked in red for display purposes. MOI: multiplicity of infection.

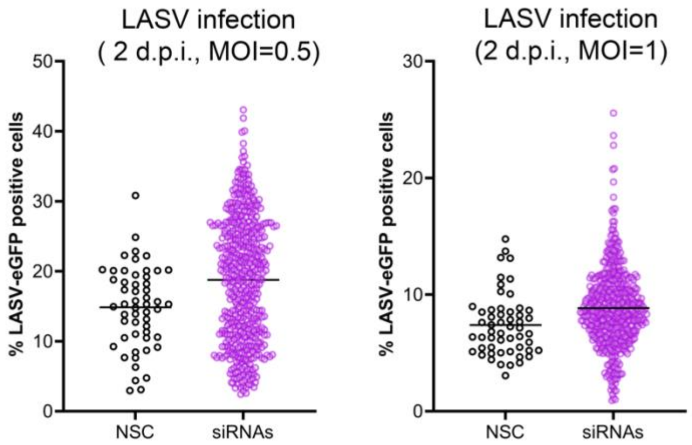


**Fig. S3. Raw infection rates of LASV-eGFP in Huh7 cells measured in siRNA screens.** The absolute percentage of cells infected with either LASV in the siRNA screen was quantified using the Cellinsight CX5 High Content Screening Platform. To visualize the range of LASV infection rate in Huh7 cells transfected with siRNA, the quantified value for the percentage infection corresponding to each well transfected with either non-silencing control (NSC) or any gene-targeting siRNA was pooled and plotted for each siRNA screen. MOI: multiplicity of infection; d.p.i.: day post infection.

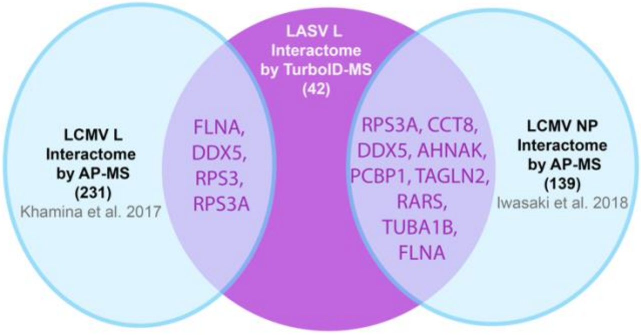


**Fig. S4. Venn diagram comparing cellular interactors of LASV polymerase with previously reported interactors of LCMV proteins.** Shared cellular proteins between LASV L polymerase interactome and LCMV interactomes are labeled in purple. AP-MS: affinity purification-based mass spectrometry.

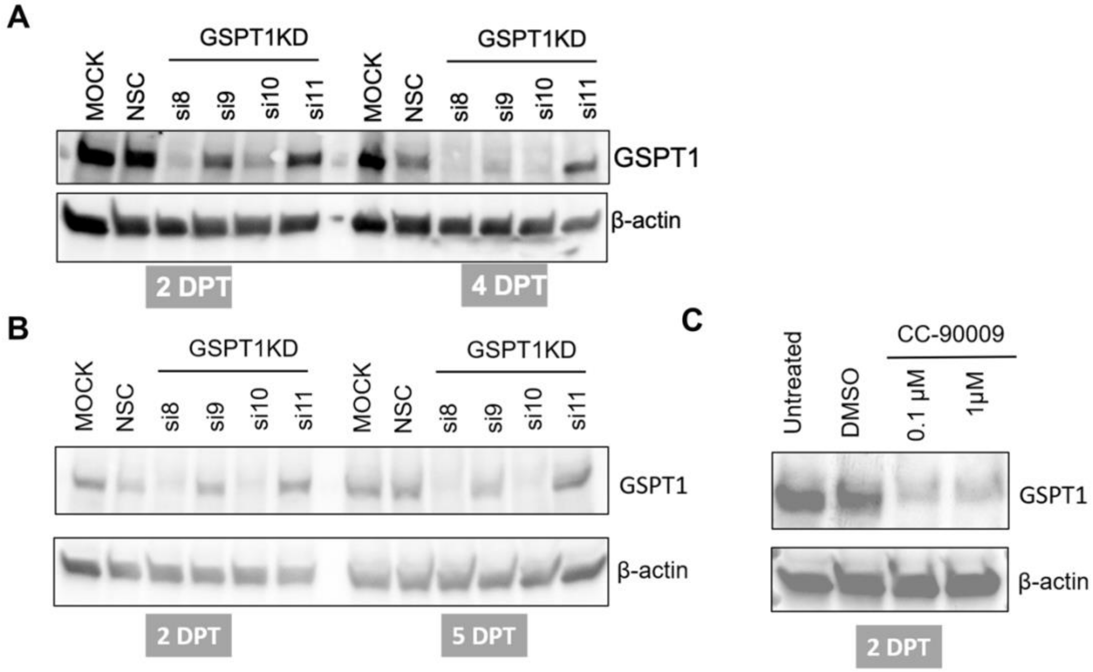


**Fig. S5. Validation of GSPT1 knockdown by siRNAs or CC-90009.** (**A, B**) Levels of endogenous GSPT1 protein (long isoform) determined with western-blotting of Huh7 cell lysates (A) or HEK 293T cell lysates (B) upon individual siRNA-knockdown from two to five days post transfection (DPT). Controls included non-transfected cells (MOCK) and cells transfected with non-silencing control (NSC) siRNA. Four individual siRNAs (si8 -10) were evaluated independently with si8 selected for GSPT1 knockdown (GSPT1 KD). **(C)** Levels of endogenous GSPT1 protein (long isoform) determined by western blotting of Huh7 cells lysates after a two-day treatment with 0.1 or 1 μM CC-90009. Controls included untreated and DMSO-treated cells. For each lane, whole cell lysates from triplicate wells were pooled for analysis.

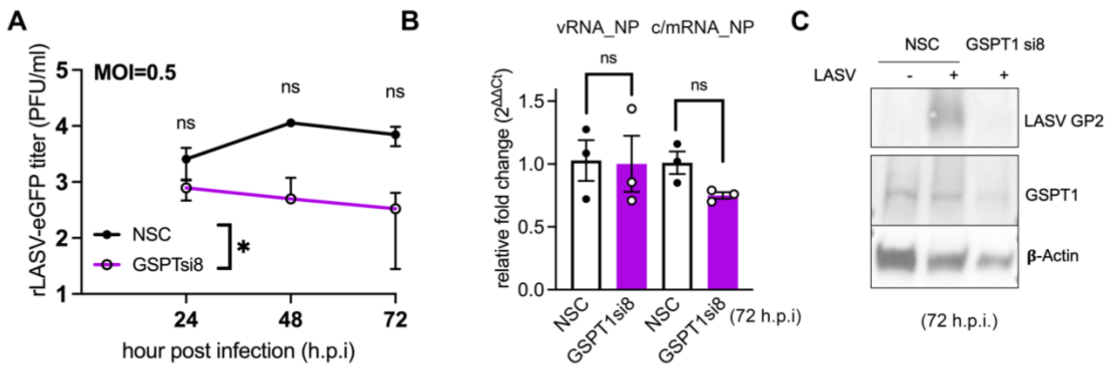


**Fig. S6. GSPT1 supports LASV multiplication in Huh7 cells. (A)** A biological replicate for the experiment that examined effects of GSPT1-knockdown (KD) on LASV viral growth kinetics in Huh7 cells. Viral titers from one experiment with technical triplicates were log-transformed and plotted as mean ± SD (error bars). Two-way ANOVA analysis with Šídák's multiple comparisons test on log-transformed titers was performed to determine the statistical significance of the effect of GSPT1-KD on viral growth kinetics and on viral titers at each time point, respectively (ns, not significant; *, $p < 0.05$). **(B)** Effect of GSPT1-KD on LASV vRNA and c/mRNA accumulation. Three individual data points from one experiment with technical triplicates are displayed and the mean ± SEM (error bars) is shown. Welch's t test was performed to determine the statistical significance of the effect of GSPT1-KD on LASV RNA accumulation. **(C)** Western blot analysis of endogenous GSPT1 protein and LASV GP2 levels in lysates from LASV-infected Huh7 cells. β-actin was used as the loading control. Lysates from triplicate wells in the growth kinetic experiments were pooled and analyzed.

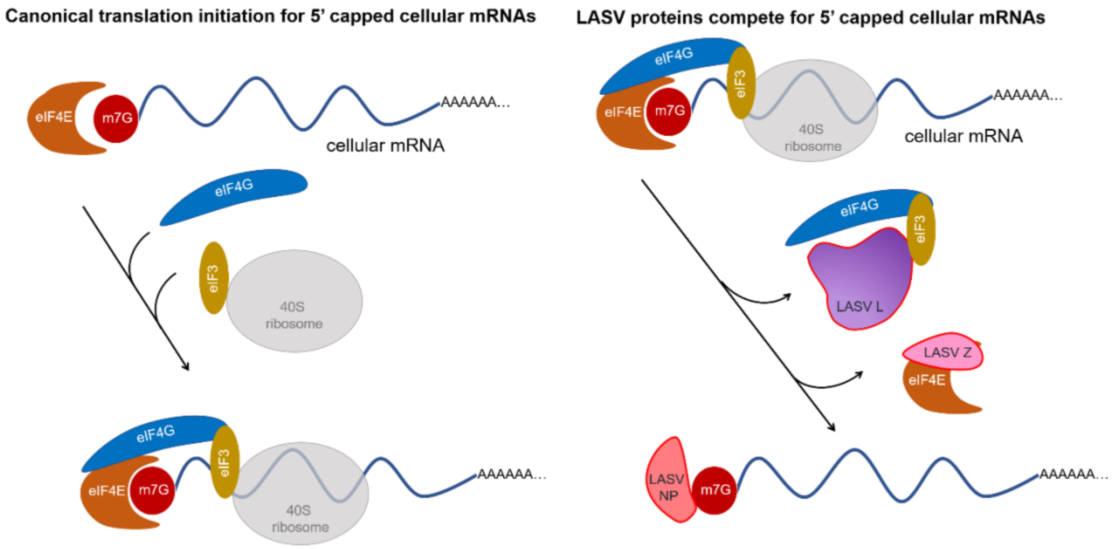


**Fig. S7. Model for competition of LASV proteins with host translational initiation factors for 5' capped mRNAs.** As part of the canonical cap-dependent translation of cellular mRNAs, the high-affinity, cytoplasmic cap-binding protein, EIF4E binds to the 5' cap structure of cellular mRNAs. Interactions between EIF4E and the 5' cap structure on cellular mRNAs are enhanced by EIF4G binding to EIF4E. EIF4G further recruits EIF3, which associates with the 40S small ribosomal subunit and other initiation factors (not shown) to form the 43S pre-initiation complex. By bridging EIF4E and EIF3, EIF4G promotes attachment of the 43S complex to the mRNA. In this model, LASV L polymerase hijacks EIF4G and EIF3, with LASV Z binding to EIF4E to reduce its cap-binding affinity. As a result, LASV NP or other proviral cap-binding proteins can retrieve the 5' cap structure to prime LASV transcription.

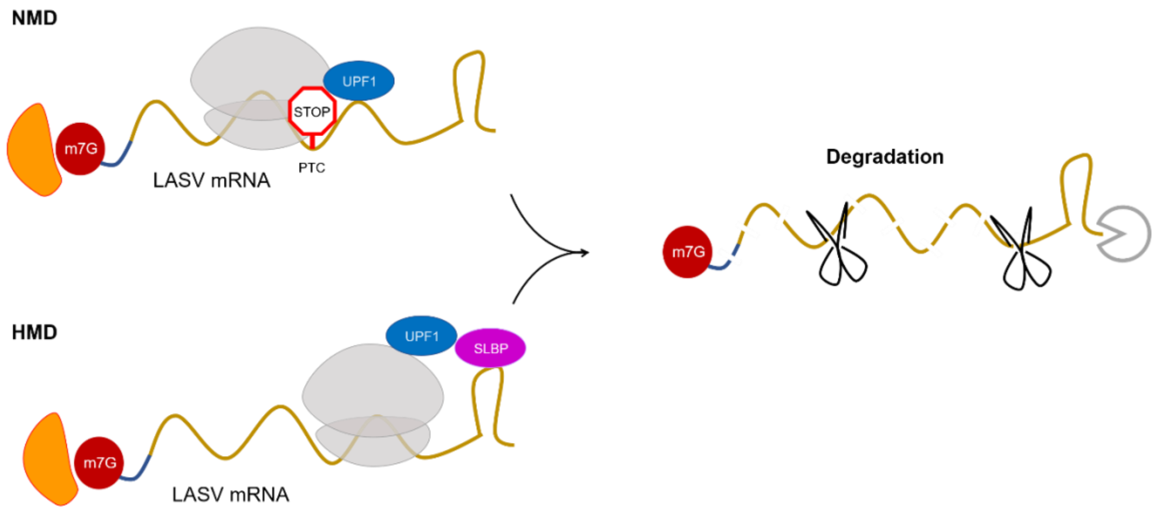


**Fig. S8. Model for LASV mRNA triggering of host RNA decay pathways.** LASV mRNAs may trigger the cellular nonsense mediated mRNA decay (NMD) pathway or the histone mRNA decay (HMD) pathway by presenting either a premature termination codon (PTC) or a 3' stem-loop structure in the mRNA.

**Dataset S1. Proteomic results.**

Relative abundance ratio and the corresponding p-value for each protein identified in all proximity-proteomic samples.

**Dataset S2. siRNA sequences.**

Sequence information for each siRNA used in the siRNA screen.

### Supplementary materials and methods

**Cell cultures and viruses:** Huh7 human hepatocytes and HEK 293T human embryonic kidney cells were maintained in Dulbecco's modified Eagle medium (DMEM-GlutaMAX) supplemented with 4.5 g/L D-Glucose, 10% fetal bovine serum (FBS), penicillin (100 U/ml), streptomycin (100 µg/ml). For viral infection experiments, Huh7 cells were cultured in Dulbecco's modified Eagle medium (DMEM) supplemented with 10% FBS and 50 µg/ml gentamicin sulfate; Vero-E6 cells were cultured in Minimum Essential Medium (MEM) supplemented with 10% FBS, 1% MEM non-essential amino acid solution, 1% sodium pyruvate solution, and 50 µg/ml gentamicin sulfate. All cells were grown at 37 °C and 5% $CO_2$.

LASV strain Josiah expressing eGFP (1) were propagated in Vero-E6 cells with 700,000 plaque forming units (PFUs/ml) of LASV-eGFP per T225 flask (at 90% confluency). LASV-eGFP was harvested at 4 days post-infection (d.p.i.). All experiments using infectious LASV were performed under biosafety level 4 (BSL-4) conditions at the Galveston National Laboratory. Virus inactivation was performed according to standard operating procedures.

LCMV Armstrong strain clone 13 variant expressing eGFP (rLCMV-eGFP) (2) was propagated by infecting BKH-21 cells at MOI of 0.01, and collecting tissue culture supernatants (TCS) at 60 hours post infection. TCS were clarified by centrifugation at 5,000 rpm for 10 minutes at 4 °C to remove cell debris. Aliquots of TCS were stored at -80 °C. Virus titers were determined using a focus forming unit assay with Vero-E6 cells.

**LASV minigenome (MG) assay:** HEK 293T cells (2 × 105 per well) were seeded in 24-well plates one day prior to transfection with the following plasmids: 300 ng pCAGGS-T7 polymerase, 300 ng pCAGGS-LASV MG encoding a ZsGreen fluorescent reporter, 300 ng pCAGGS-LASV-L, 150 ng pCAGGS-LASV-NP. In some cases: pCAGGS-LASV-L was replaced with a tagged variant. A control was included in every assay in which the pCAGGS-L plasmid was omitted to verify active MG RNA synthesis mediated by LASV L. Plasmids were transfected with Lipofectamine 3000 transfection reagent (1.5 µl/well). At 72 hours post-transfection, cell lysates were collected and green fluorescence intensity was measured.

**Biotinylation with TurboID fusion proteins:** HEK 293T cells seeded in 24-well plates were transfected with the LASV minigenome system including the LASVL-HA-TurboID fusion protein. As a control, HEK 293T cells were transfected with an equimolar amount of plasmid encoding an HA-tagged TurboID. Biotin (500 µM) was added to cell culture media three days post-transfection. Multiple labeling time points were sampled for optimization. The reaction was stopped by placing cells on ice. Excess biotin was removed by three times of cold DPBS wash, and soluble whole cell lysates were collected for western blot analysis using streptavidin-HRP and antibodies against HA tag or LASV NP.

**Immunofluorescent analysis (IFA) using confocal microscopy:** Acetone cleaned, glass coverslips (1.5 mm thickness) placed inside wells of 24-well plates were treated with human fibronectin (50 mg/ml) for 20 min in a 37 °C incubator, prior to seeding of HEK 293T cells (4X104/well). Twenty-four hours later, cells in the monolayer were transfected with the LASV minigenome (MG) system using Trans-IT LT1 as described. For in-cell biotinylation experiment, biotin (500 µM final concentration) was added to the culture media allowing and incubated for 1 hour to allow in-cell biotinylation of cells transfected with the LASV MG components. Labeling was stopped by washing cells with cold DPBS. For all IFA specimens, cells were fixed in 4% paraformaldehyde for 15 mins at room temperature, permeabilized with 0.1% Triton-X100 for 10 mins, and blocked in 1% normal goat serum for at least one hour. Primary antibody incubations were performed at room temperature for two hours or at 4 °C overnight. Specimens were washed three times with PBS-0.1% Tween-20 and incubated with secondary antibodies for one hour at room temperature. Nuclei were counterstained with Hoechst. Confocal images were acquired on Zeiss LSM880-airyscan system under super-resolution mode, using a 63x/NA1.4 oil objective. Please see SI Appendix for full description of antibodies used in IFA experiments.

**Sample preparation for in-cell biotinylation and streptavidin enrichment:** A protocol adapted from Branon et al. (3) was used. Briefly, HEK 293T cells were seeded in 6-well plates ($1X10^6$ cells per well) one day before transient transfection using Trans-IT LT1 and plasmids expressing wild-type LASV L (L-WT) or the TurboID-fused LASV polymerase (L-HA-TurboID), both in combination with plasmids expressing LASV NP, T7 polymerase, and a T7-polymerase driven LASV minigenome. To enable in-cell biotinylation, HEK 293T cell monolayers three days post-transfection were incubated with 500 µM biotin containing complete media for four hours. Biotinylation was stopped by washing monolayers with cold DPBS twice before collecting the cell pellets by centrifugation at 1500 rpm for 3 mins at 4 ˚C. Cell pellets (1V) were lysed in one volume of 1XRIPA buffer containing protease inhibitors (Roche) and benzonase followed by incubation on ice for 15-20 mins. Soluble lysates were clarified by centrifugation at 13000 rpm for 5 mins at 4 ˚C.

For streptavidin enrichment, streptavidin-magnetic beads (80 µl slurry/mg of total protein) were washed with 1X RIPA buffer twice and incubated with 2-3 mg soluble cell lysate for one hour at room temperature. The beads were then washed twice with 1 ml RIPA buffer, once with 1 ml 1M KCL, once with 1 ml 0.1M $Na_2CO_3$, once with 1 ml 2 M Urea in 10 mM Tris-HCl (pH 8.0), and twice with 1 ml RIPA lysis buffer. The washed beads were then either eluted in 50 µl SDS loading buffer supplemented with 20 mM DTT and 2 mM biotin for quality checks or processed for on-bead digestion. Quality checks for successful streptavidin enrichment included silver staining and western blotting of SDS-PAGE-separated input and streptavidin-enriched samples side-by-side.

For on-bead digestion, proteins bound to beads were washed twice with 50 mM Tris HCL (pH 7.5) and twice with 2 M urea/50 mM Tris buffer (pH 7.5). A final volume of 80 µl of 2 M urea/50 mM Tris containing 1 mM DTT and 0.4 µg trypsin was added to the washed beads, which were incubated overnight with shaking at 37 ˚C. Digested supernatants containing biotinylated peptides were transferred to a new tube. The streptavidin beads were washed twice with 60 µl 2 M urea/50 mM Tris buffer (pH 7.5) and the remainder was combined with the on-bead digest supernatant. The eluate was reduced with 4 mM DTT for 30 mins at room temperature with shaking, and then alkylated with 10 mM IAA (iodoacetamide) for 45 mins in the dark at 25 ˚C with shaking. An additional 0.5 µg of trypsin was added to the sample and the digestion was completed overnight with shaking at 37 ˚C. After a final digestion, samples were acidified by adding formic acid to a final concentration of 1 % and stored at -80 ºC before shipment to the Scripps Florida Proteomic core for further processing, labeling (for TMT labeling), and mass-spectrometry analysis.

**Proximity proteomics (with tandem-mass-tag labeling):** Following on-bead trypsin digestion of biotinylated proteins, Lassa (WT_pol and TurboID_pol) samples were acidified by the addition of 1% formic acid, desalted with 10 µg-capacity C18 StageTips (Thermo Fisher Scientific, Waltham, MA), and dried under vacuum. Peptides were resuspended in 100mM TEAB, labelled with TMT labels (6-plex) according to the manufacturer's instructions (Thermo Fisher Scientific, Waltham, MA) and pooled.

The labelling scheme for three biological replicates of each condition was as follows:

| LASSA sample | WT_pol_01 | WT_pol_02 | WT_pol_03 | TurboID_pol_01 | TurboID_pol_02 | TurboID_pol_03 |
|---|---|---|---|---|---|---|
| TMT 6-plexs label | 126 | 127 | 128 | 129 | 130 | 131 |

The pooled and multiplexed samples were dried under vacuum, re-solubilized in 1% TFA and then desalted using 2 µg-capacity C18 ZipTips (Millipore, Billerica, MA) before drying again under a vacuum. For mass spectrometry, dried TMT-labelled peptides were reconstituted in 5 µL 0.1% TFA, vortexed briefly, and then sonicated for 15 min. The peptides were subsequently on-line eluted into a Fusion Tribrid mass spectrometer (Thermo Fisher Scientific, San Jose, CA) from an Acclaim PepMapTM RSLC C18 nano Viper analytical column (2 mm, 100 Å, 75-µm ID × 50 cm, Thermo Scientific, San Jose, CA) using a gradient of 5-25% solvent B (80/20 acetonitrile/water, 0.1% formic acid) in 180 mins, followed by 25-44% solvent B in 60 mins, 44-80% solvent B in 0.1 min, a 5-mins hold of 80% solvent B, a return to 5% solvent B in 0.1 min, and finally a 20 min hold of solvent B. All flow rates were 300 nL/min delivered using a nEasy-LC1000 nano liquid chromatography system (Thermo Fisher Scientific, San Jose, CA). Solvent A consisted of water and 0.1% formic acid. Ions were created at 2.1-2.5 kV using the EASY-Spray™ ion source held at 50 ˚C (Thermo Fisher Scientific, San Jose, CA). A synchronous precursor selection (SPS)-MS3 mass spectrometry method based on the work of Ting et al. (4), was used with scanning between 380-2000 m/z at a resolution of 120,000 for MS1 in the Orbitrap mass analyzer, and performing CID at top speed in the linear ion trap of peptide monoisotopic ions with charge 2-8, using a quadrupole isolation of 0.7 m/z and a CID energy of 35%. The top 10 MS2 ions in the ion trap between 400-1200 m/z were then chosen for HCD at 65% energy and detection in the Orbitrap at a resolution of 60,000 and an AGC target of 1E5. The injection time was 120 msec (MS3). Three technical replicates were performed for all six biological samples.

**Bioinformatic analysis of mass spectrometry data:** The raw data for technical replicates were added as "fractions" into Proteome Discover v 2.3 (Thermo Fisher Scientific, San Jose, CA) and quantitative analysis of the TMT experiments was performed simultaneously with protein identification. The precursor and fragment ion mass tolerances were set to 10 ppm, 0.6 Da, respectively; the enzyme was trypsin with a maximum of 2 missed cleavages. Uniprot Human proteome FASTA files with added sequences specific for LASSA viral and TurboID fusion proteins was used in SEQUEST searches. Impurity correction factors obtained from Thermo Fisher Scientific for each kit were included in the search and quantification. The following settings were used to search the streptavidin-enriched data. Dynamic modifications: Oxidation, +15.995Da (M); Deamidated, +0.984 Da (N, Q); TMT6plex, +229.163 Da (K); and Biotin, +226.078 Da (K). Static modifications: TMT6plex, +229.163 Da (N-Terminus) and Carbamidomethyl, +57.021 (C). Only unique +Razor peptides were considered for quantification purposes. The Target Decoy feature of Proteome Discoverer 2.3 was used to set a false discovery rate (FDR) of 0.01. Total peptides quantified were normalized to Streptavidin to adjust for loading bias and the Protein Abundance Based method was used to calculate the protein level ratios. The Low Abundance Resampling method was used to impute missing data and co-isolation threshold and SPS Mass Matches thresholds were set to 50 and 65, respectively. ANOVA (Individual Proteins) was performed using Proteome Discoverer 2.3 workflow and a FDR of 0.05 was chosen as the cut off to identify the top tier proteins that are differentially expressed across WT and TurboID samples in the LASSA experiment. Sequences added to the Uniprot Human proteome database for this experiment were Nucleoprotein [Lassa mammarenavirus] and LASV_L_TurboID_internal fusion protein.

**Thresholding and network analysis of LASV polymerase-proximity interactome:** To curate a list of high-confidence hits for the LASV polymerase-proximity interactome, we applied a threshold of " > 1" for the $\log_2$ transformed relative abundance ratio (TurboID_pol/WT_pol) and a threshold of "< 0.05" for the adjusted *p-value* for each abundance ratio. The list of high-confidence hits in the LASV polymerase interactome, including critical parameters used in thresholding, is presented in **Dataset S1**. The LASV polymerase interactome composed of high-confidence proteomic hits was visualized by Cytoscape. The web-based bioinformatic analyzer STRING was used to perform functional enrichment analysis (FDR < 1%) to categorize nodes within the LASV polymerase interactome based on the biological processes each node is involved in.

**siRNA functional screening and data analysis:** Huh7 cells (5 x $10^3$ per well) were seeded in black 96-wells plate having a clear bottom; edge wells were filled with DPBS. Twenty-four hours, Huh7 monolayers were transfected with 10 nM of individual siRNAs using Lipofectamine RNAiMAX (0.1 μl per well). Knock down (KD) of each target was performed using four individual siRNA transfections, each one performed in triplicate. Each plate included two sets of triplicated internal control wells transfected with 10 nM All Star negative control (NSC) to account for plate-to-plate variations. Forty-eight hours after siRNA transfection, plates were transferred to BSL4 for infection with rLASV-eGFP (1) [S segment: GenBank #MH358389], [L segment: GenBank #MH358388] at the indicated MOI. Infected monolayers were fixed and inactivated with 10% formalin at 48 hours post infection and removed from the BSL4 facility. The percentage of LASV-eGFP infected Huh7 cells (% infection) with different siRNA treatments were quantified with the Cellinsight CX5 High Content Screening Platform using a 10x objective. The cell count in each well was determined by the number of Hoechst-stained nuclei.

The relative percentage of virus infection and relative cell count were calculated by normalizing raw percent viral infection rate and cell count of individual siRNA treatment to corresponding control wells in each plate. Heat maps showing relative %virus infection and relative cell count were generated by GraphPad Prism 9 using the normalized value of each individual siRNA treatment. The mean of numerical values from each triplicate well are used for plotting. To quantitatively analyze the impact of siRNA treatment on viral infection and cell count, multiple t-tests were performed to compare the normalized percentage infection for each siRNA treatment to each corresponding NSC control and determined the P value or each comparison.

Bubble plots were generated by plotting numerical values of normalized cell count for each siRNA treatment against the value for the normalized percentage of infection for data points with $p \leq 0.05$ (P value represented by the size of each scatter point). Outliers having a cell count that differed significantly from the average by one standard deviation were ruled out because the effect of siRNA treatment on percentage of infection in these cases can be confounded by a marked change in the total number of cells. siRNAs used for the screen were from the Genome-wide ON TARGET-Plus (OTP) Human siRNA library from Dharmacon/Thermo Scientific. Sequence information for each siRNA been used is shown in **Dataset S2**.

**LASV growth kinetics experiment:** Huh7 cells ($5 \times 10^4$ per well) were seeded in 24-well plates and transfected with 10 nM siRNA (NSC or GSPT1 si8, experimentally validated) the next day as described above. Forty-eight hours after transfection, Huh7 monolayers were transferred into BSL-4 facilities and infected with rLASV-eGFP at MOI of 0.5 (PFU/cell) for one hour. Infection of the cells was followed by removal of the inoculum and replenishing with 2% FBS containing fresh media. For drug treatment, rLASV-eGFP infected Huh7 cells (MOI=0.5) were incubated either with CC-90009 at the specified concentration or DMSO (vehicle) post infection. For virus titration, aliquots of supernatant were harvested at the indicated time points from infected monolayers and used to infect Vero-E6 cells for foci-quantification. At indicated time points, infected Huh7 monolayers were either fixed with 10% neutral buffered formalin and stained with DAPI for fluorescent imaging, or collected directly in Trizol or 4x Laemmli buffer for inactivation and downstream RNA/protein quantifications, Equal volumes of Laemmli lysates were loaded on SDS-PAGE gels for western blot analysis. Equal amounts of total RNA in each sample were used for strand-specific RT-qPCR.

**LCMV growth kinetics experiment:** Huh7 cells ($1.5 \times 10^5$ per well) were seeded in 24-well plates, and infected with rLCMV/GFP-P2A-NP [S segment: GenBank # DQ361065] [L segment: GenBank #DQ361066] (2) at MOI of 0.5 (PFU/ml) for 1.5 hour. Cell infection was followed after removing the inoculum and replenishing with complete media containing 10% FBS and either CC-90009 at the specified concentration or DMSO (vehicle). For virus titration, aliquots of supernatant were harvested at the indicated time points from infected monolayers and used to infect Vero-E6 cells for foci-quantification. At the indicated time points, infected Huh7 monolayers were either fixed with 4% paraformaldehyde (PFA) and stained with Hoechst, or collected directly in Trizol or 2x protein loading buffer for downstream RNA/protein quantifications. Equal volumes of whole cell lysates were loaded on SDS-PAGE gel for western blot analysis. Equal amounts of total RNA in each sample were used for RT-qPCR.

**Co-Immunoprecipitations (co-IP):** FLAG-tag-based, co-IP reactions were performed as previously described (5). HEK 293T cells ($1X10^6$ per well) were seeded in 6-well plates 24 hours before transfection of cells in each well with 500 ng of pCI-empty or pCI-FLAG-GSPT1 plasmid combined with 2 µg plasmid expressing HA-tagged LASV L protein. For some conditions, 1 µg LASV NP plasmid, 2 µg T7-polymerase driven minigenome plasmid and 2 µg T7 polymerase plasmid were included in the co-transfection mix. Three days later, the transfected cells were washed with DPBS and lysed in 1X RIPA buffer including protease inhibitor cocktail (cOmplete™, EDTA-free Protease Inhibitor Cocktail), on ice for 15 mins. Soluble whole cell lysates were clarified by centrifugation at 12,000X rpm for 10 min. FLAG-M2 affinity beads (40 µl slurry per reaction) were washed with DPBS once and twice with NT2 buffer before blocking with BSA in NT2 buffer (1 mg/ml) for 30-60 min at room temperature. Lysates containing equal amounts of total protein (1-1.5 mg determined by BCA assay) were incubated with BSA-blocked FLAG-M2 affinity beads and incubated overnight at 4 ˚C with rotation. Supernatant containing unbound proteins was removed by centrifugation at 5000 x g for 30 s, followed by four washes in 1 ml NT2 buffer. Immunoprecipitated proteins were eluted in 4X-SDS-loading buffer containing 10% beta-mercaptoethanol following by 10 min incubation at 95 ˚C before western blotting analysis.

For HA-tag based, co-IP reactions, HEK 293T cells ($1X10^6$ per well) were seeded in 6-well plates. Cells in each well were transfected 24 hr later with 2.5 µg plasmid expressing HA-tagged LASV L protein or HA-tagged GFP-Halotag fusion protein. Three days later, transfected cells were washed with DPBS and lysed in 1X RIPA buffer including protease inhibitor cocktail (cOmplete™, EDTA-free Protease Inhibitor Cocktail), and benzonase nuclease, on ice for 15 mins. Soluble whole cell lysates were processed and subjected to HA-IP described in the FLAG-IP section except with anti-HA affinity matrix.

**RT-qPCR:** To quantify LASV RNA from infected cells, total RNA was extracted from Trizol-inactivated samples in each well using Direct-zol RNA Miniprep Plus according to the manufacturer's protocol and eluted in 50 µl DEPC-treated water. RNA was quantified using a Nanodrop spectrophotometer. Total RNA (200 ng) from each sample was used as the template for reverse transcription (RT) using a High-Capacity cDNA Reverse Transcription Kit according to the manufacturer's protocol. Strand-specific RT primers binding to the LASV NP gene were used to amplify vRNA or c/mRNA for the LASV S segment. Resulting cDNAs were diluted 1:100 in DEPC-treated water and then used for quantitative real-time PCR with a Bio-Rad CFX Real-Time System. SYBR select master mix was added to the diluted cDNA template and target specific qPCR primer pairs in a 20 µl reaction volume. Each reaction was performed in triplicate wells on qPCR plates. PCR conditions were those for the Standard Cycling Mode (Primer Tm ≥ 60 ˚C) according to the manufacturer's protocol. A default dissociation curve was performed immediately after the real-time PCR to obtain the Tm (melting temperature) of each target. The Tm of samples using the same qPCR primer pairs were confirmed to be the same. Amplification plots have the baseline subtracted, and the relative quantification (ΔΔCT) method was used to analyze results. GAPDH was used as the house-keeping gene for data normalization. Samples from NSC or DMSO were used as a control to calculate the relative fold-change of the target amplicon in GSPT1-knockdown (GSPT1si8) or CC-90009 treated samples.

To quantify LCMV RNA from infected cells, total RNA was extracted from Trizol-solubilized samples in each well using a column free method. The extracted RNA was resuspended in 12 µl 1 mM sodium citrate, pH 6.5 ± 0.1 buffer. Total RNA (500 ng) from each sample was used as the template for reverse transcription (RT) using Superscript IV Reverse Transcriptase according to the manufacturer's protocol. Random hexamers were used to amplify total RNAs. The resulting cDNAs (10 ng) were used in the quantitative real-time PCR using PowerUp SYBR Green Master Mix and LCMV NP-specific qPCR primers. Each reaction was performed in triplicate wells of qPCR plates. PCR condition was following the Standard Cycling Mode (Primer Tm ≥ 60 ºC) according to manufacturer's protocol. Amplification plots have the baseline subtracted, and the relative quantification (ΔΔCT) method was used to analyze results. GAPDH was used as the house-keeping gene for data normalization. Samples from DMSO treated cells were used as a control to calculate the relative fold-change of the target amplicon in CC-90009 treated samples.

**Cell viability quantification by Cell-titer Glo assay:** Confluent Huh7 cells seeded in 96-well plates were incubated with cell culture media containing DMSO or CC-90009 . Cell viability upon drug treatment was measured at the indicated time points using cell-titer Glo 2.0 (Promega). The percentage cell viability of each drug treatment was calculated by normalizing luminescence measured in drug-treated cell lysates to that of DMSO-treated cell lysates.

Figures 2A, 3A, 5A, 6D and S1 were created using BioRender.

**Table S1.** Plasmids used in this work.

| Plasmid name | Description | Resource |
|---|---|---|
| pCAGGS-LASV-NP | Encodes wild-type NP of Lassa virus | de la Torre Lab (6) |
| pCAGGS-LASV-L | Encodes wild-type L protein of Lassa virus | de la Torre Lab (6) |
| pCAGGS-LASV-L-HA | Encodes N-terminal HA-tagged L protein of Lassa virus | de la Torre Lab (6) |
| pCAGGS-LASV-L-TurboID | Encodes modified Lassa L protein with an internal HA-tag and the full length of TurboID L protein (insertion site:407/408, SRIT/Q). TurboID sequence was amplified from the plasmid 3XHA-TurboID-NLS_pCDNA3 as a gift from Alice Ting (3).) | Engineered for this work |
| pT7-LASV-ZsGreen-MG | Encodes a ZsGreen reporter with regulatory sequences from authentic Lassa genome (s segment) | de la Torre Lab (6) |
| pCAGGS-T7pol | Encodes a cytoplasmic-located, bacteriophage T7 RNA polymerase | de la Torre Lab (6) |
| pCI-empty | Does not encode any perceivable protein, serve as transfection control plasmid | Engineered for this work |
| pCI-FLAG-GSPT1 | Encodes a N-terminal FLAG-tagged GSPT1/eRF3a (long isoform, 68.7KDa) amplified from and modified based on the plasmid (pCI-MS2V5-eRF3a F76a) as a gift from Niels Gebring (7). | Engineered for this work |
| pCAGGS-eGFP | Encodes an eGFP reporter | de la Torre Lab |
| pCAGGS-empty | Does not encode any perceivable protein, serve as transfection control plasmid | de la Torre Lab |
| pCDNA5-HA-eGFP-Halotag2 | Encodes an HA-tagged eGFP-Halotag2 fusion protein, served as control for HA-IP | A kind gift from Craig Crews (8) (Addgene plasmid # 41742 ; http://n2t.net/addgene:41742 ; RRID:Addgene_41742) |

**Table S2.** Primers used in this work.

| Primer name | Sequence (5' to 3') | Used for | Target |
|---|---|---|---|
| RT-LASV-vRNA | GTGGACACAATCTTTGAGGAGG | Reverse transcription (RT) | LASV vRNA (s segment) |
| RT-LASV-c/mRNA | TCACAGAACGACTCTAGGTG | RT | LASV c/mRNA (s segment) |
| Oligo dT (20mer) | TTTTTTTTTTTTTTTTTTTT | RT | Polyadenylated mRNAs |
| LASV_NP_F | GGAATGAGTGGTGGTAATCAAGG | qPCR | LASV NP coding sequence |
| LASV_NP_R | TTTTCACATCCCAAACTCTCACC | qPCR | LASV NP coding sequence |
| Random Hexamers | N/A | RT | Total RNAs |
| LCMV_NP_F | CAGAAATGTTGATGCTGGACTGC | qPCR | LCMV NP coding sequence |
| LCMV_NP_R | CAGACCTTGGCTTGCTTTACACAG | qPCR | LCMV NP coding sequence |

**Table S3.** Antibodies and compound used in this work.

| Antibodies and affinity-conjugate | Source | Resource | Used in |
|---|---|---|---|
| Streptavidin-HRP | N/A | Thermo Fisher, Invitrogen (Cat#SA10001) | WB (1:1000) |
| Anti-HA | Mouse monoclonal (16B12) | Biolegend (Cat#901513) | WB (1:1000), IF (1:500) |
| Anti-HA | Rabbit monoclonal (C29F4) | Cell signaling technology (Cat#3724) | WB (1:1000) |
| Anti-β-actin | Mouse monoclonal (AC-15) | SantaCruz Biotech (Cat#sc-69879) | WB (1:4000) |
| Anti-LASV NP | Mouse monoclonal (61SP) | Zalgen/Autoimmune Technology | WB (1:1000) |
| Anti-LCMV GP | Mouse monoclonal (G204) | de la Torre Lab | WB (1:500) |
| Anti-LASV GP | Mouse-human chimeric (MHC-13.4E) | A kind gift from James E. Robinson (9) | WB (1:500) |
| Anti-GSPT1/eRF3 | Rabbit polyclonal | Abcam (Cat#ab49878) | WB (1:1000), IF (1:500) |
| Anti-PSMC5 | Rabbit monoclonal [EPR13565(B)] | Abcam (Cat#ab178681) | WB (1:1000) |
| Anti-RPS3 | Rabbit monoclonal (EPR7808) | Abcam (Cat#ab128995) | WB (1:2000) |
| Anti-EIF4G2 | Rabbit monoclonal (D88B6) | Cell Signaling Technology (Cat#5169) | WB (1:1000) |
| Anti-RARS | Rabbit polyclonal | Biorbyt (Cat# orb247357) A kind gift from Paul Schimmel | WB (1:5000) |
| Anti-AIMP2/JTV1 | Mouse monoclonal (1D6B5) | Proteintech (Cat#66848) A kind gift from Paul Schimmel | WB (1:2000) |
| Anti-UPF1 | Rabbit monoclonal (D15G6) | Cell Signaling Technology (Cat#12040) | WB (1:1000) |
| Streptavidin-AlexaFluor594 | N/A | Thermo Fisher, Invitrogen (Cat#s32356) | IF (1:500) |
| Anti-mouse-HRP | goat | Thermo Fisher, Invitrogen (Cat#31437) | WB (1:1000) |
| Anti-rabbit-HRP | goat | Southern Biotech (Cat#4050-05) | WB (1:1000) |
| Anti-mouse-AlexaFluor568 | goat | Thermo Fisher, Invitrogen (Cat#A11004) | IF (1:500) |
| Anti-mouse-AlexaFluor647 | goat | Thermo Fisher, Invitrogen (Cat#A21236) | IF (1:500) |
| Anti-rabbit-AlexaFluor647 | goat | Thermo Fisher, Invitrogen (Cat#A27040) | IF (1:500) |

| | | | |
|---|---|---|---|
| Streptavidin-magnetic beads | N/A | Thermo Fisher (Cat#88817) | Streptavidin pull-down, Proteomic sample prep |
| Anti-FLAG M2 affinity Gel | Mouse monoclonal (M2) conjugated to agarose beads | Sigma-Aldrich (Cat# A2220) | FLAG-IPs |
| Anti-HA affinity matrix | Rat monoclonal (3F10) covalently coupled to agarose beads | Roche (Cat# 11815016001) | HA-IPs |
| CC-90009 | N/A | MedKoo Biosciences (Cat# 207005) | LASV/LCMV growth kinetics experiment (0.1/ 1$\mu$M) |
